# Deciphering Bedaquiline and Clofazimine Resistance in Tuberculosis: An Evolutionary Medicine Approach

**DOI:** 10.1101/2021.03.19.436148

**Authors:** Lindsay Sonnenkalb, Joshua Carter, Andrea Spitaleri, Zamin Iqbal, Martin Hunt, Kerri Malone, Christian Utpatel, Daniela Maria Cirillo, Camilla Rodrigues, Kayzad S. Nilgiriwala, the CRyPTIC Consortium, Philip W. Fowler, Matthias Merker, Stefan Niemann

## Abstract

Bedaquiline (BDQ) and clofazimine (CFZ) are core drugs for treatment of multidrug resistant tuberculosis (MDR-TB), however, our understanding of the resistance mechanisms for these drugs is sparse which is hampering rapid molecular diagnostics. To address this, we employed a unique approach using experimental evolution, protein modelling, genome sequencing, and minimum inhibitory concentration data combined with genomes from a global strain collection of over 14,151 *Mycobacterium tuberculosis* complex isolates and an extensive literature review. Overall, 230 genomic variants causing elevated BDQ and/or CFZ MICs could be discerned, with 201 (87.4%) variants affecting the transcriptional repressor (Rv0678) of an efflux system (mmpS5-mmpL5). Structural modelling of Rv0678 suggests four major mechanisms that confer resistance: impairment of DNA binding, reduction in protein stability, disruption of protein dimerization, and alteration in affinity for its fatty acid ligand. These modelling and experimental techniques will improve personalized medicine in an impending drug resistant era.

## Introduction

Multidrug-resistant (MDR: resistance to at least isoniazid [INH] and rifampicin [RMP]) Mycobacterium tuberculosis complex (Mtbc) strains represent a serious challenge for global tuberculosis (TB) control^1,2^. The World Health Organization (WHO) estimates that close to half a million people worldwide were infected in 2019 with an RMP-resistant (RR) Mtbc strain, of whom 78% were MDR^1^.

Compared to patients with sensitive TB, treating MDR-TB treatment takes longer (years), the drugs are less effective and more toxic, and cure rates are low (about 60% globally)^1,3^. As a consequence, ineffective MDR-TB regimens in many high incidence settings have led to an expansion of drug resistant Mtbc strains. The WHO has recommended new treatment regimens for MDR-TB, including new and repurposed drugs such as bedaquiline (BDQ) and clofazimine (CFZ)^4–7^. Both drugs are now central to MDR-TB therapy, and are also part of the recently WHO-endorsed shorter, all-oral MDR-TB regimen^7,8^.

Selection experiments have found resistance to BDQ and or CFZ as mediated by mutations in the genes *atpE, pepQ, Rv1979c*^9–11^, and *Rv0678* – the latter of which seems to be the most clinically relevant and strains with BDQ associated mutations in *Rv0678* and *atpE* have been selected in multiple patients^12–17^. Rv0678 is a marR-like transcriptional regulator of the mmpS5-mmpL5 efflux pump with a canonical N-terminal winged helix-turn-helix (wHTH) DNA binding domain and a C-terminal helical dimerization domain. The Rv0678 protein forms a homodimer which binds to the promotor region of the mmpS5-mmpL5 operon^18–21^. This repression is reversed upon the conformational change of the Rv0678 molecule in the presence of a fatty acid ligand^20^. Importantly, Mtbc strains with mutations in *Rv0678* have been shown to confer resistance against both BDQ and CFZ^22,23^, thus undermining the recently WHO-endorsed treatment guidelines^8^. Moreover, mutations in *Rv0678* are the most common resistance mechanism to CFZ and BDQ found in patient isolates today^16,17,24–29^.

So far, mutations scattered across the full gene length of *Rv0678* (498bp) have been found to occur *in vitro* and in patients isolates leading to variable shifts in BDQ and CFZ minimum inhibitory concentrations (MICs)^13,29,30^. However, a comprehensive understanding of the associations between *Rv0678* mutations and their resulting phenotype, as well as their structural effects on the transcriptional repressor Rv0678, is lacking.

In this paper we present the results of an experimental model that evolves Mtbc strains under sub-lethal drug concentrations. Such a weak selection pressure has been shown to select for diverse pheno- and genotypes in other bacterial species and likely also reflects the physiological conditions in TB patients^31–33^. Recent studies indicate the inability of most drugs to diffuse into the granuloma at therapeutic levels, including CFZ and BDQ which strongly bind caseum macromolecules^34–38^. BDQ has further been shown to accumulate in cellular compartments of various types of macrophages (including neutrophils), but was unevenly distributed in intracellular bacteria^39,40^. Single resistant clones and total bacterial populations from consecutive cultivation passages were selected at clinically relevant drug concentrations, analyzed by whole genome sequencing (WGS), and compared with mutations identified in 14,151 patient isolates collected by the CRyPTIC consortium. Selected mutations were further investigated computationally for their putative effects on the Rv0678 protein structure and function, which provides a starting point for design of computational algorithms to predict drug resistance. The data generated provide a comprehensive resistance mutation catalogue for genotypic BDQ/CFZ drug susceptibility testing and will improve MDR-TB treatment design and likely outcomes for patients receiving a BDQ and/or CFZ containing therapies.

## Results

### Experimental evolution under sub-lethal drug concentrations

To better understand the resistance mechanisms of Mtbc strains against BDQ (which often includes correlated resistance to CFZ), we evolved Mtbc strains under sub-lethal drug concentrations. The experimental set-up was developed to allow for a high number of bacterial generations under a moderate drug selection pressure, thus increasing the mutation diversity compared to classical mutant selection experiments, i.e. exposing a culture at a late exponential phase to the critical drug concentrations (Figure S1). Our H37Rv laboratory strain (ATCC 27294) MIC was 0.25 mg/L for BDQ and 0.5 mg/L for CFZ, in high bacterial density liquid culture (10^7^ CFU/mL inoculum) – we shall refer to these as the wild-type (WT) MICs. During five culture passages spread over 20 days, we exposed H37Rv to either BDQ or CFZ at one of four concentrations, ranging from one-half to one-sixteenth of the WT MIC (Figure S2), and selected colonies on solid media (7H11 plates) supplemented with 0.12 mg/L (1:2 MIC) and 0.25 mg/L (MIC) for BDQ and 0.25 mg/L (1:2 MIC) for CFZ. At passages 1, 3, and 5, single colonies as well as the whole bacterial population were genotypically characterized, and single selected mutations were phenotypically defined (elevated MIC) (Tables 1, S1).

**Table 1:**
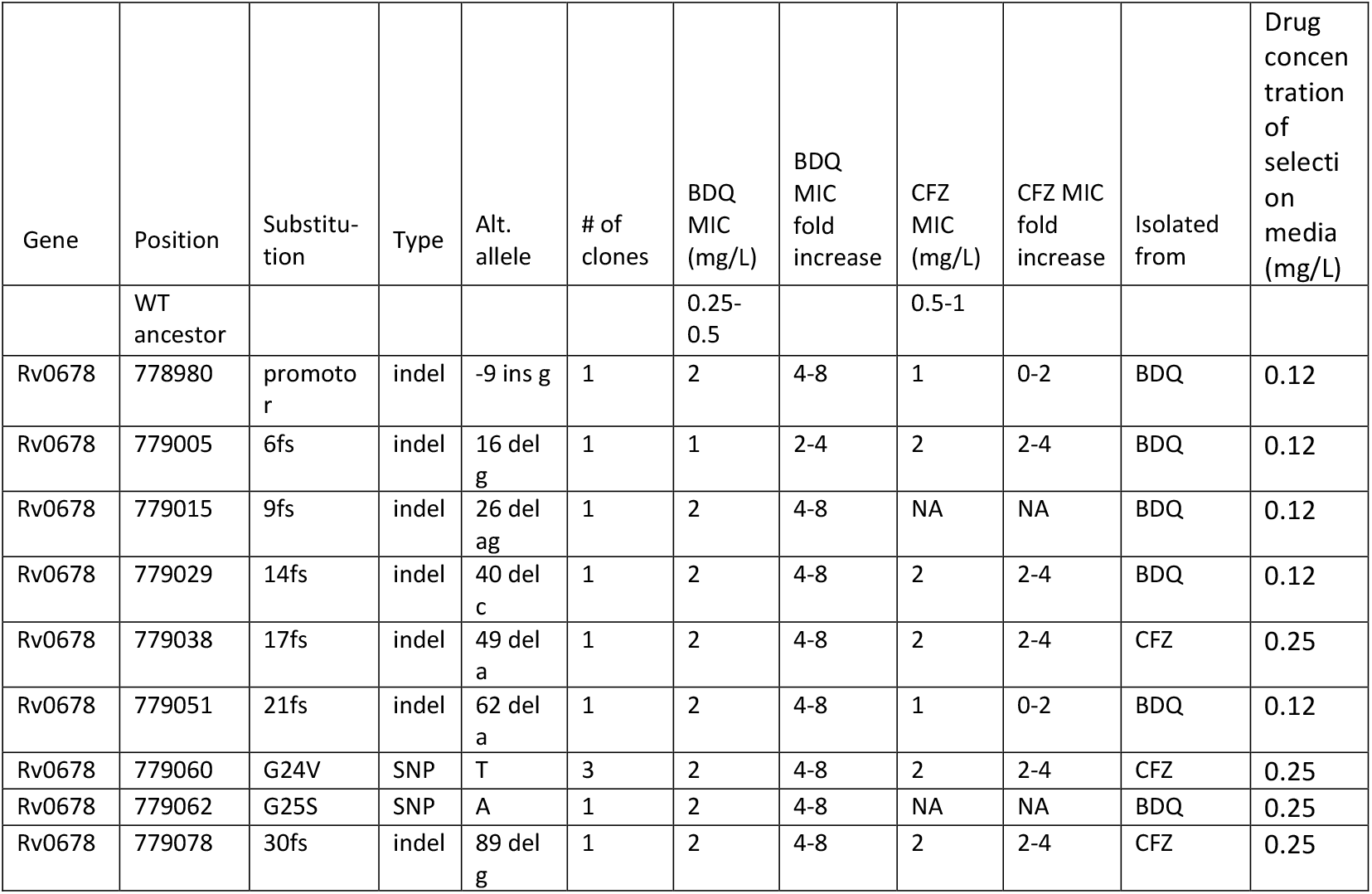
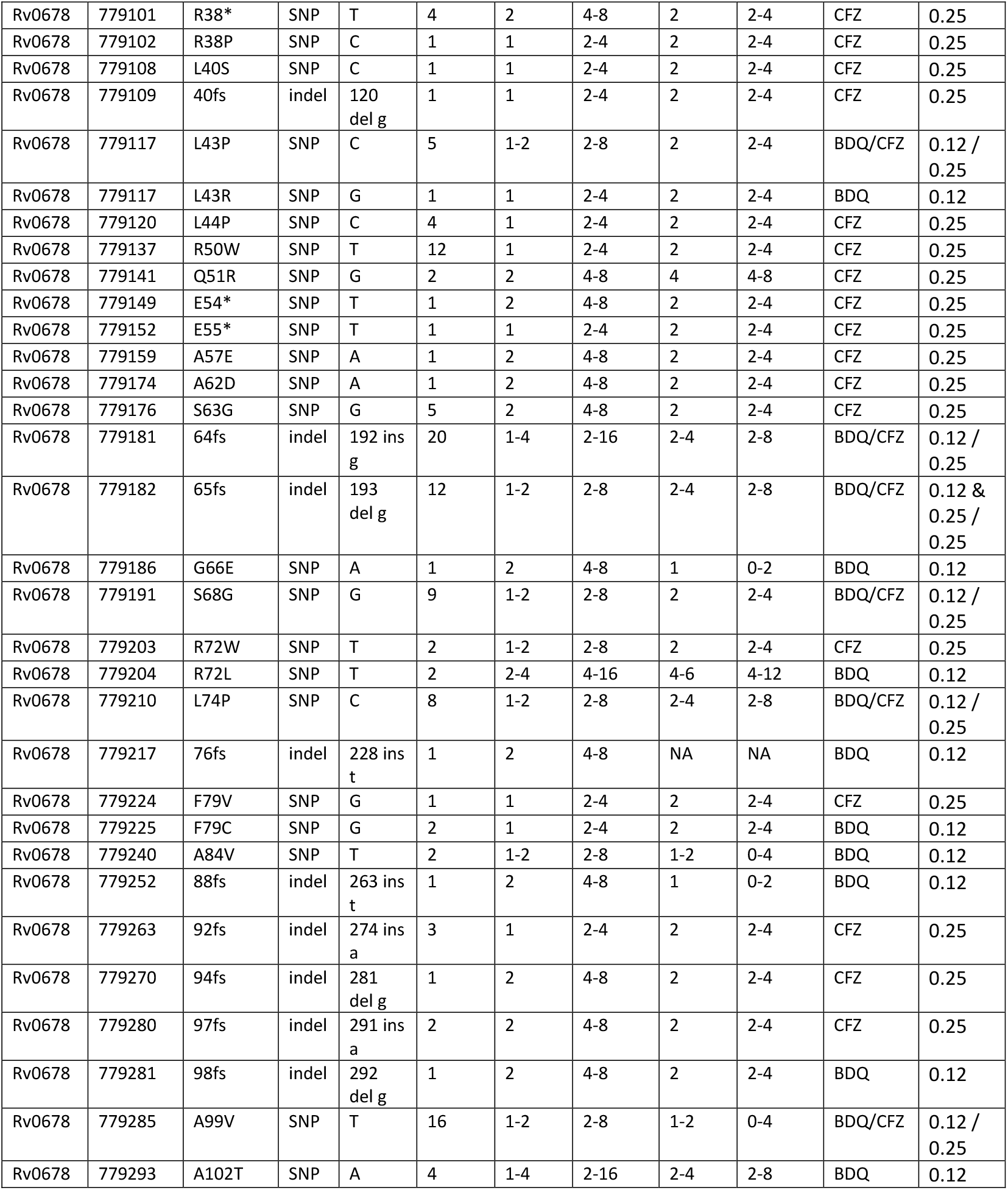
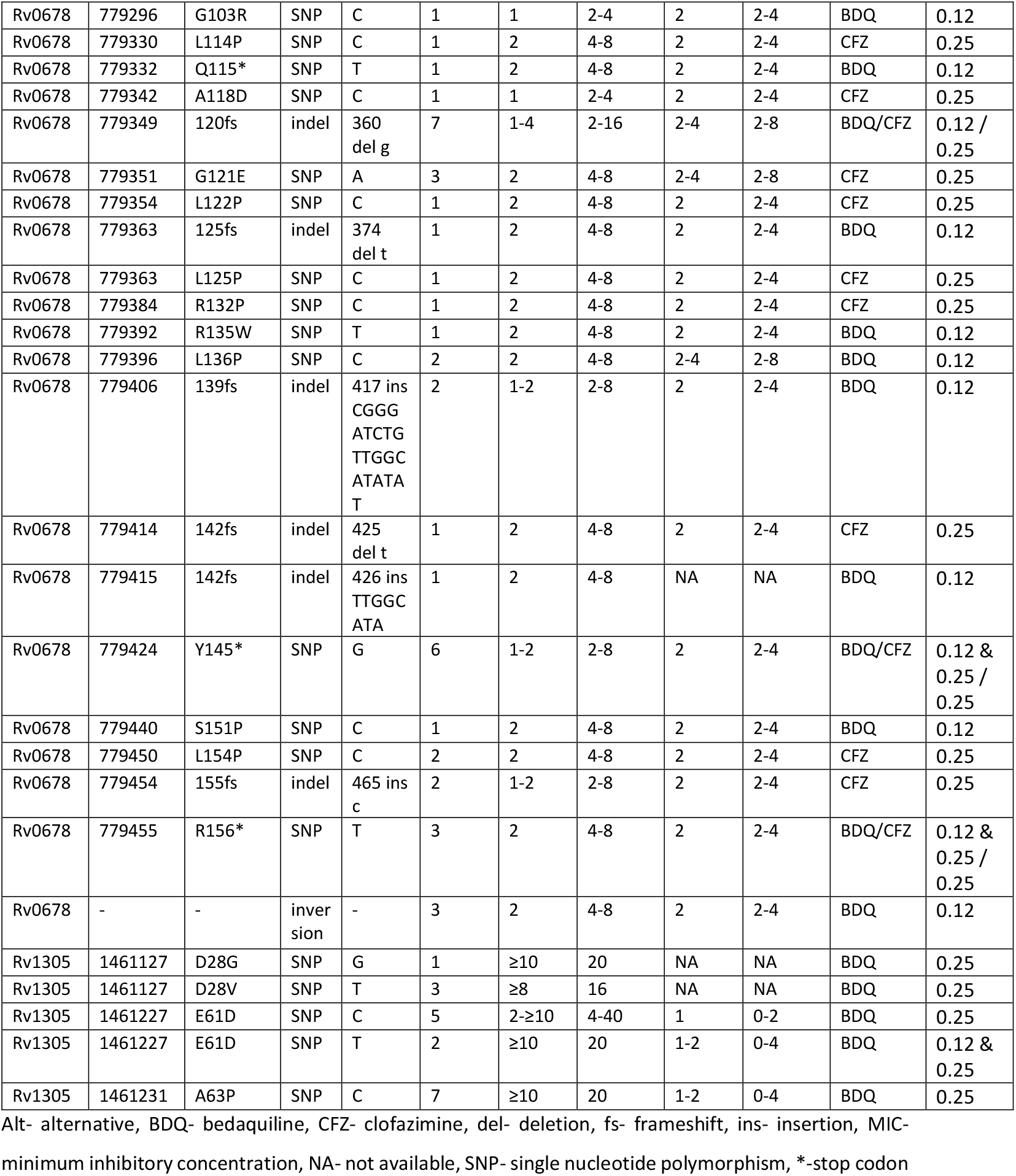
*In vitro* selected mutations conferring BDQ resistance. Genotype of each variant was determined by whole genome sequencing, and substitution annotated by coding position, “alternative allele” denotes base pair change. Minimum inhibitory concentration (MIC) was determined by resazurin microtiter plate assay and compared to wild type (WT) susceptible ancestor to describe MIC fold increase. The number of times a mutation was independently selected was included under “# of clones”. Finally, from which experiment (BDQ or CFZ) the mutant was isolated from included as “isolate from”.

Strikingly, exposure to BDQ or CFZ concentrations as low as one-eighth of the WT MIC over 20 days (12 to 19 generations) led to the selection and enrichment of significantly more resistant cells as compared to growth in the absence of the drugs (P=2.8 x 10^-10^, and P=2.4 x 10^-3^, respectively, Figure S2). As expected, there was less enrichment of resistant populations at early timepoints, e.g. after 4 days.

### Phenotypic and genotypic characterization of BDQ resistant mutants

In total, we randomly selected 270 single colonies from BDQ- and CFZ-supplemented agar plates that had been inoculated with samples from passages 1 (4 days), 3 (12 days), and 5 (20 days), of two independent experiments (Figure S1). In total, 203/270 mutants were successfully pheno- and genotyped (67 were excluded due to DNA library preparation issues or inadequate growth). The single colonies were sub-cultured and the MICs for BDQ and CFZ were determined for each clone using broth microtiter dilution. All clones exhibited an elevated MIC at least 2-fold higher than the WT drug susceptible ancestor and we identified in total 61 unique variants with one mutation in the *Rv0678* gene per clone. We additionally selected five unique mutations in the *atpE* (*Rv1305*) gene (Table 1).

Clones which harbored a mutation in *Rv0678* had a 2-4-fold MIC increase for BDQ, whereas mutations in *atpE*, exhibited 4 to over 10 times the MIC of the WT strain (Table 1). For 12 randomly selected clones with seven different variants we verified the results obtained in broth microtiter dilution plates using a Mycobacterium growth indicator tube (MGIT) assay. These experiments showed comparable MICs of the mutant clones with those obtained from the previous method (Table S2).

The 61 unique mutations in *Rv0678* affected 54 different codons (Table 1), out of which 21 (34%) were frameshift (fs) mutations, 34 (56%) had one non-synonymous SNP, and 6 (10%) a premature inserted stop codon. The mutations were scattered over the entire sequence of *Rv0678* and no dominant cluster could be identified (Figure 1A). However, the four mutations which were most frequently observed were: 192 ins g (64fs), A99V, 193 del g (65fs), and R50W; these mutations were detected in 19, 17, 13, and 12 single selected clones, respectively (Table 1). Notably, nucleotides 192-198 were also identified as a hotspot for frameshifting variation in a prior review, potentially due to the homopolymeric nature of this region^41^.

**Figure 1:**
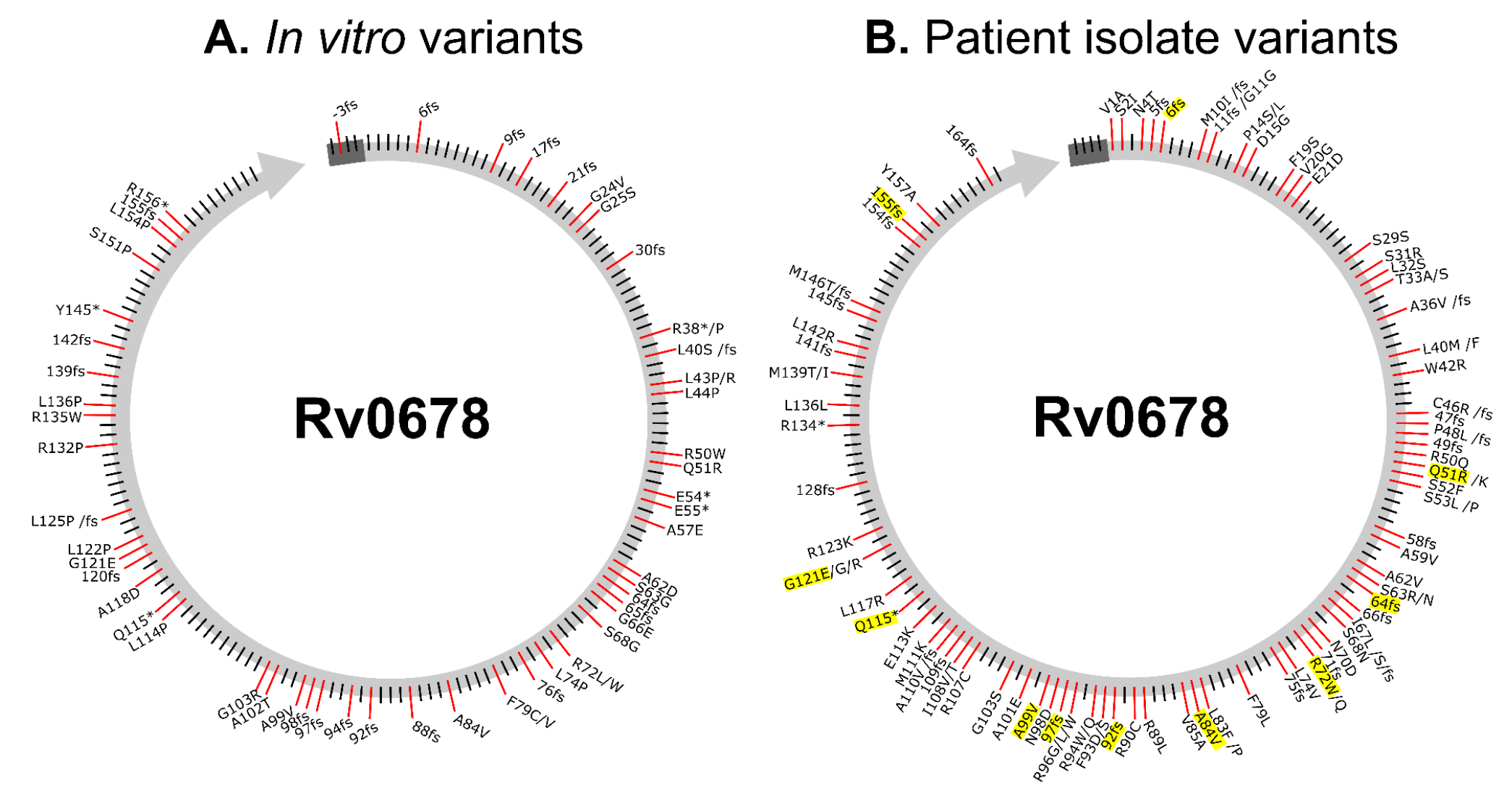
BDQ resistance associated variants mapped in *Rv0678* gene. Resistance associated mutations detected within patient dataset and *in vitro* selection were designated along Rv0678 gene at corresponding coding positions by non-synonymous single nucleotide polymorphisms, and by a protein disrupting mutation such as a stop codon insertion (*) or frameshift (fs) mutations. Data sets divided by (A) mutations which were selected in *in vitro* experiments with elevated minimum inhibitory concentrations verified to bedaquiline, and (B) mutations which were detected in patient isolates either collected by CRyPTIC or described in published literature. Overlapping mutations in observed in both patients and *in vitro* are highlighted in yellow.

As mentioned before, no mutant clone had more than one resistance mediating variant in *Rv0678* or *atpE*, however, 38 mutants harbored a second mutation in one of 14 genes that have not been previously proposed to be associated with resistance against BDQ and/or CFZ (Table S3). The most common mutation found in addition to *Rv0678* mutations was R119H in *Rv1890c* – this gene codes for a conserved hypothetical protein^42^, and the mutation was selected 11 times along with six different resistance mutations. The second most common co-selected mutation was *130R in *Rv1871c* (a conserved hypothetical membrane protein^43^) that was selected seven times along with four different resistance mutations. Other secondary mutations occurred in genes which were involved in cell wall synthesis, information pathways, metabolism/respiration, protein regulation, or lipid metabolism (Table S3).

In addition to single colony sequencing, we also employed a more unbiased approach based on population sequencing of all the mutant colonies on a given plate, in order to elucidate all possible variants. On average the genome wide coverage for a total of 81 samples was 308±50. We then reported mutations if the position was covered by at least one read in both forward and reverse orientations and there were two reads with a phred score over 20. This identified 45 additional mutations in *Rv0678* and one in *atpE* (A63T), as well as four mutations in *pepQ* (*Rv2535c*; F97V, G96G, V92G, A87G) (Table S1). As in the single colony analysis, we found the following mutations dominating in different independent evolutionary experiments and selected across different drug concentrations: *Rv0678* 192 ins g (64fs), A99V, R50W, and L43P.

### Resistance caused by huge inversion interruption in Rv0678

Three mutants with elevated BDQ (2 mg/L) and CFZ (2 mg/L) MICs but lacking mutations in the BDQ/CFZ resistance associated genes were further subjected to long read sequencing with the PacBio Sequel II system. A *de novo* assembly employing the PacBio SMRT^®^ Link software resulted in one closed Mtbc genome, and two assemblies with 1 and 4 four contiguous sequences (contigs), respectively. All assemblies covered more than 99.9% of the H37Rv reference genome (NC_000962.3). All three assemblies showed a large-scale sequence re-arrangement with the borders at position 779,073 (Rv0678 coding sequence) and 3,552,584 (intergenic region), flanked by a transposase open reading frame (IS*6110*). A 2.5Mb fragment was inverted and thus split the *Rv0678* gene into halves (Figure 2), which was not detected by a classical reference mapping approach and revealed a further interesting resistance mechanism. We scanned all BDQ resistant samples in the CRyPTIC strain collection for similar events using local assembly with ARIBA^44^. In one isolate we assembled one contig which clearly represented a similar inversion disrupting *Rv0678*, but since the depth of support was just 1 read (mean, across the contig) and this was Illumina (short read) data, we considered this only circumstantial evidence.

**Figure 2:**
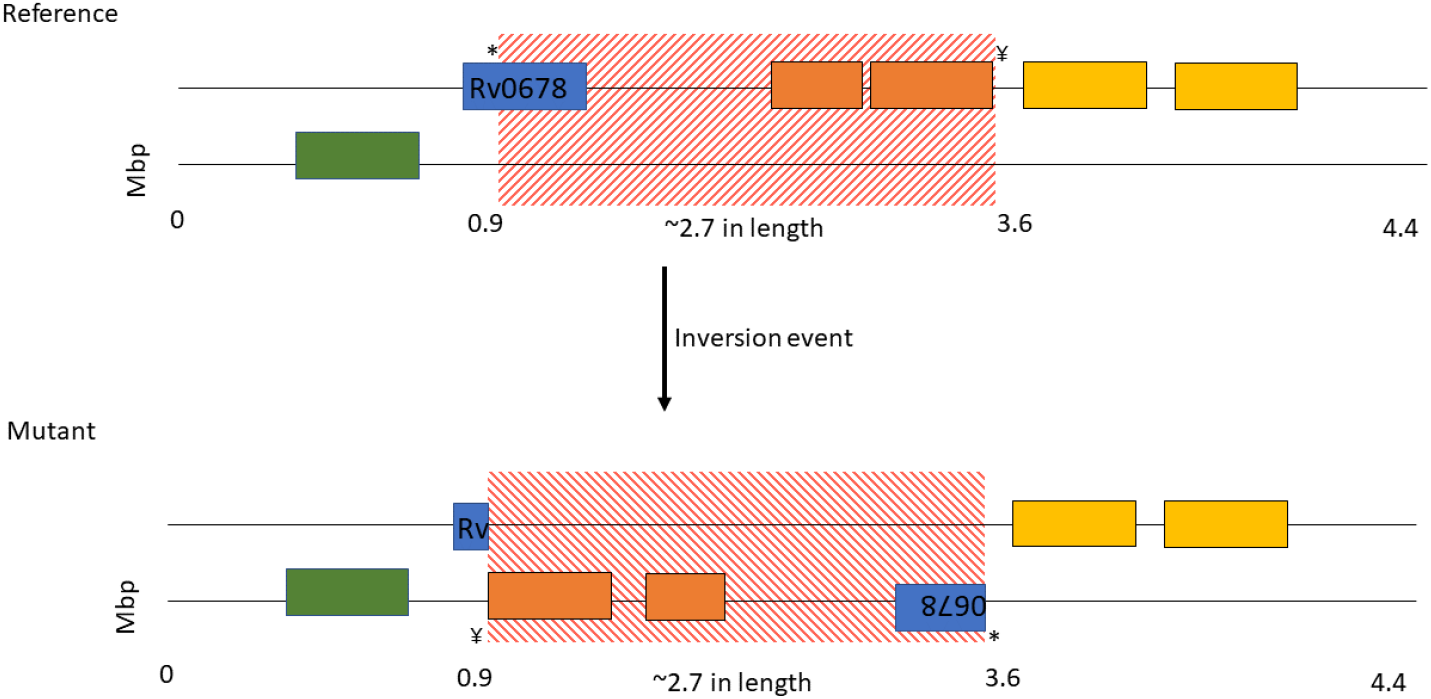
Large scale gene rearrangement in Rv0678 gene. *De novo* assembly using PacBio SMRT^®^ Link software indicated a 2.7 Mb inversion (red) impacting *Rv0678* (blue), at positions 779,073 coding sequence (amino acid 34 in Rv0678) and 3,552,584 intergenic region; flanked by transposase related genes (orange and yellow).

### Mutations associated with bedaquiline and clofazimine resistance in clinical isolates

Next, we aimed to comprehensively describe the BDQ and CFZ resistant variants occurring in patients, to develop a catalogue of known resistance variants. We searched the literature for patient reports describing BDQ resistance during MDR-TB therapy and analyzed pheno- and genotypes of 14,151 clinical Mtbc strains collected by the CRyPTIC consortium (Table S4). In this patient dataset, we identified 98 mutations throughout *Rv0678*, 71 of which were not previously described in patients (Table 2). In addition, seven variants in *Rv0678* harbored mutations found in different lineages (Table S4), pointing to convergent evolution at these sites. Eleven mutations from the patient-derived set were also shared with our *in vitro* derived datasets (selected variants and population sequencing) (Figure 1B & 3A).

**Figure 3.**
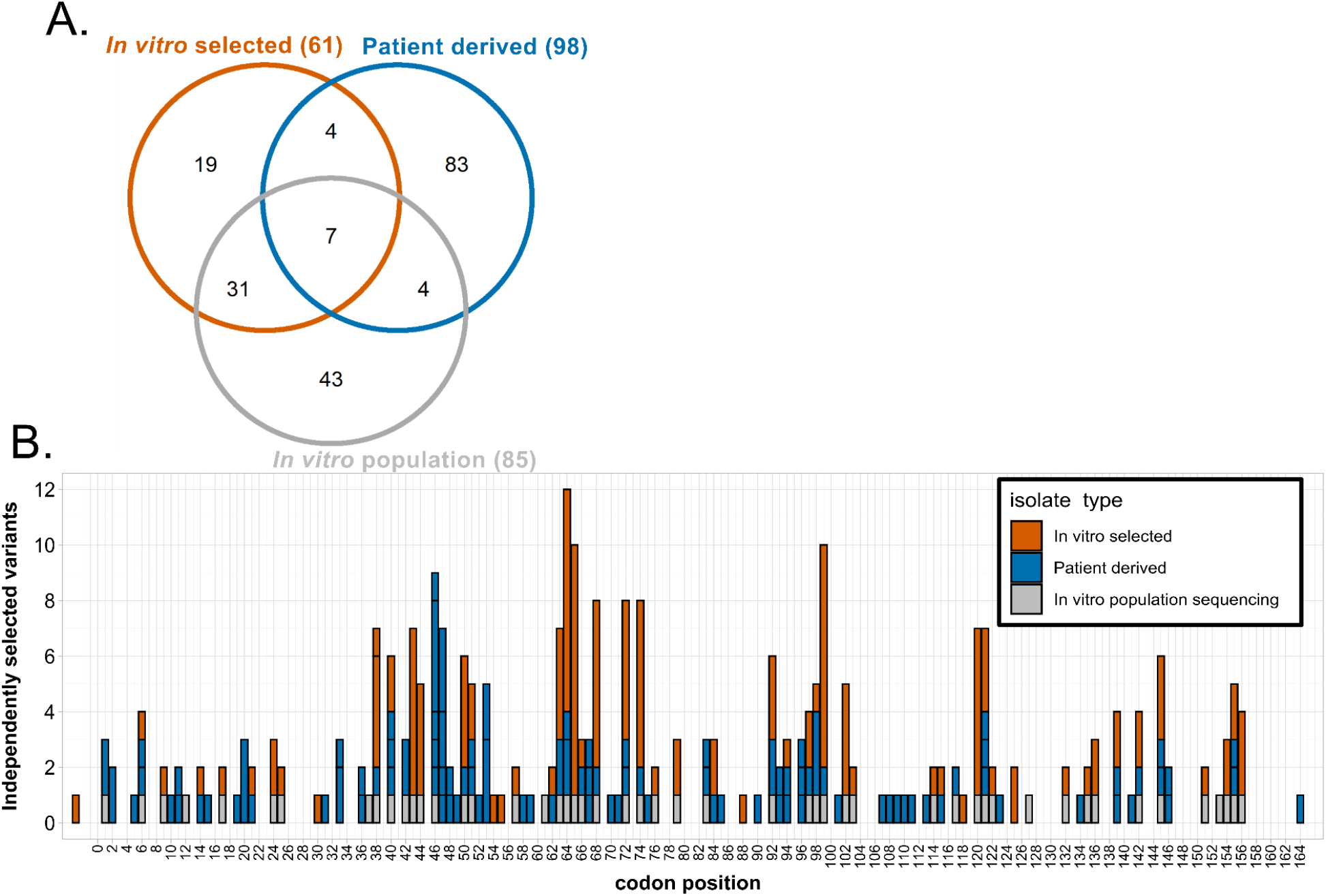
Mutations and coding position overlap throughout *Rv0678, in vitro* and patient datasets. Overlapping mutations which conferred BDQ resistance were compared between three datasets: *in vitro* selected mutants (Table 1), *in vitro* population sequencing (Table S1), and patient derived variants (Table 2). (A) Venn diagram describes number overlapping mutations in the three datasets, all mutations must have the same SNP or a deletion/insertion at same coding position to overlap. (B) Data represents number of mutants selected at given coding position for *in vitro* selected (orange) and patient derived variants (blue), indicated with gray box if detected in *in vitro* population sequencing.

**Table 2:**
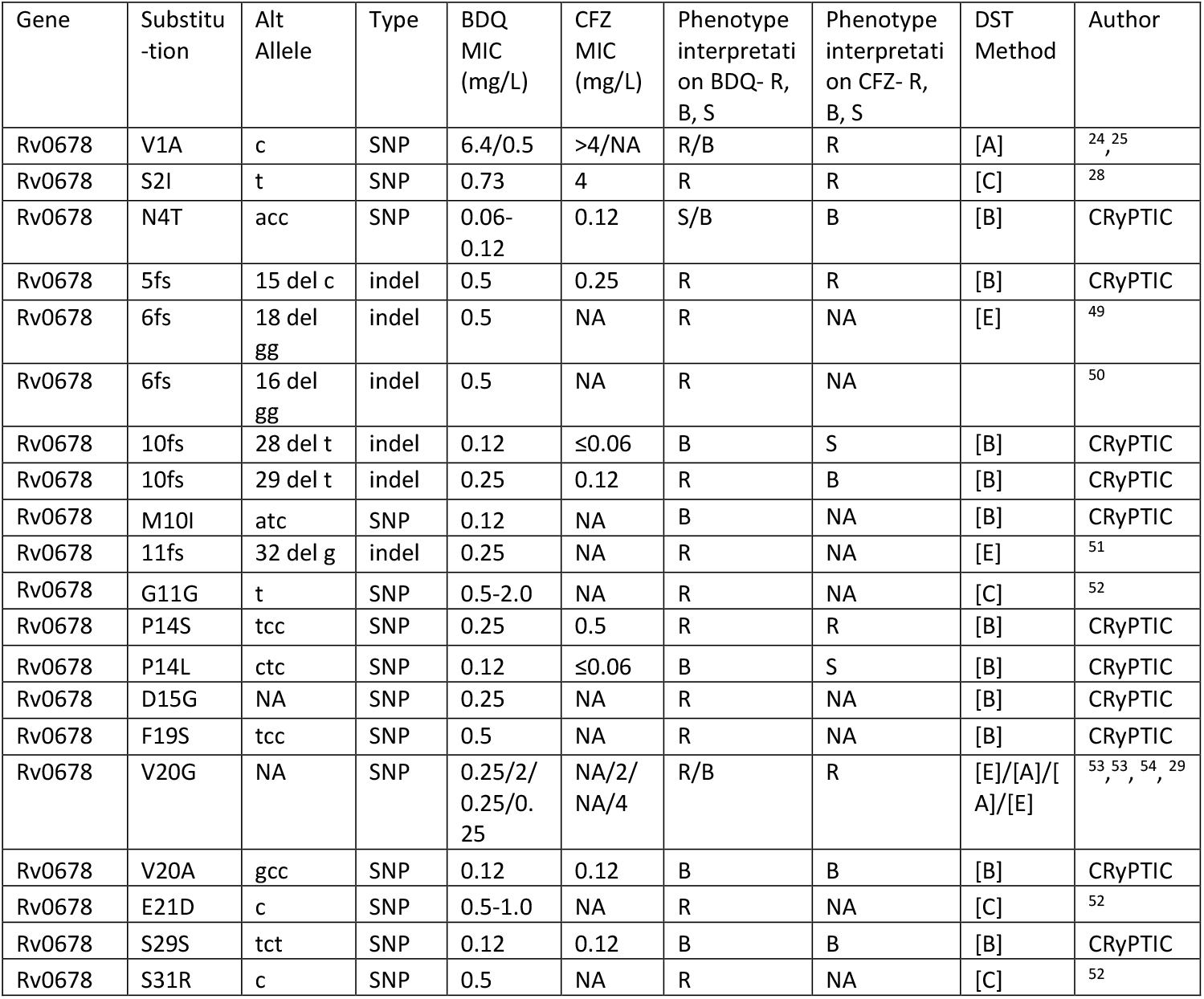
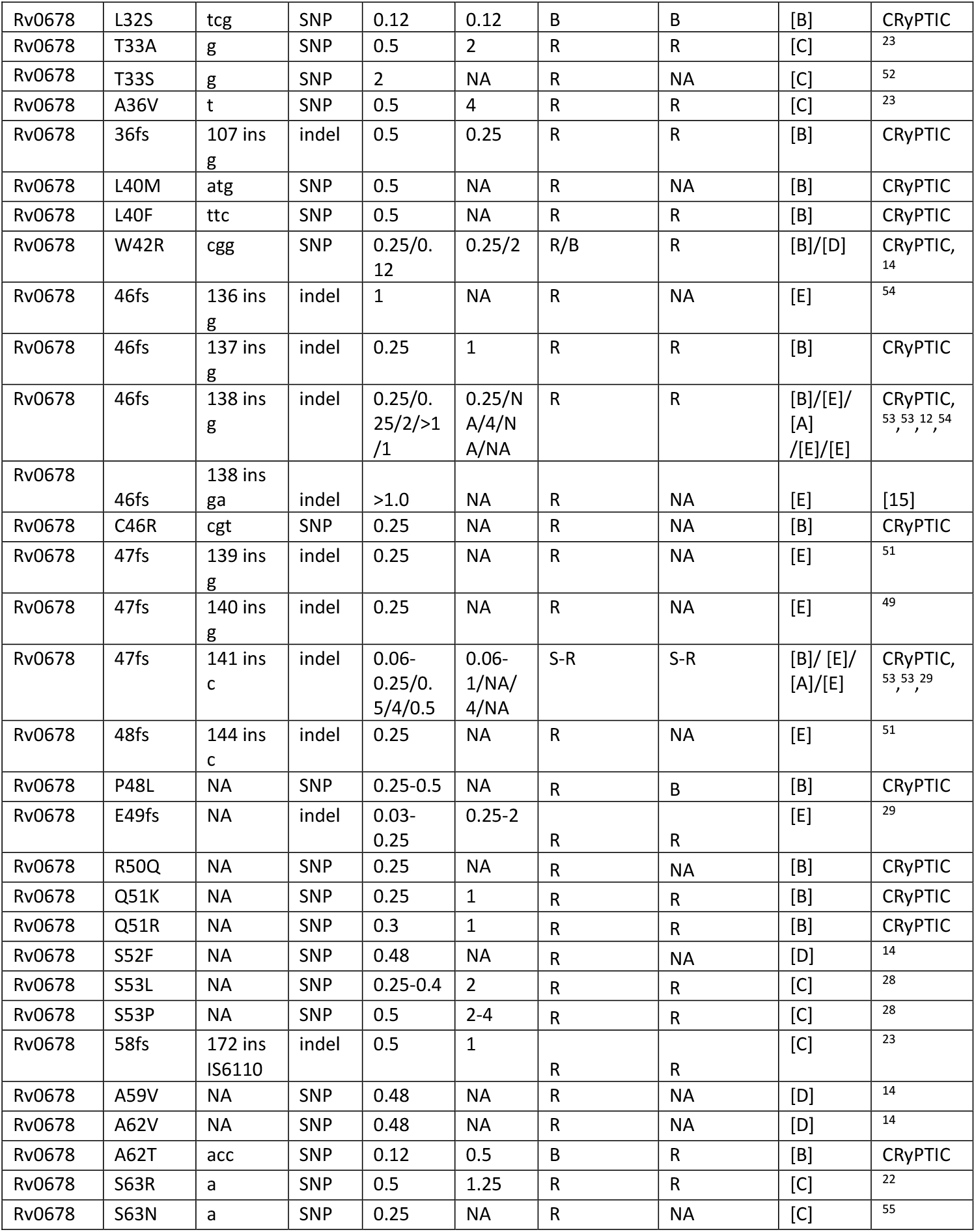
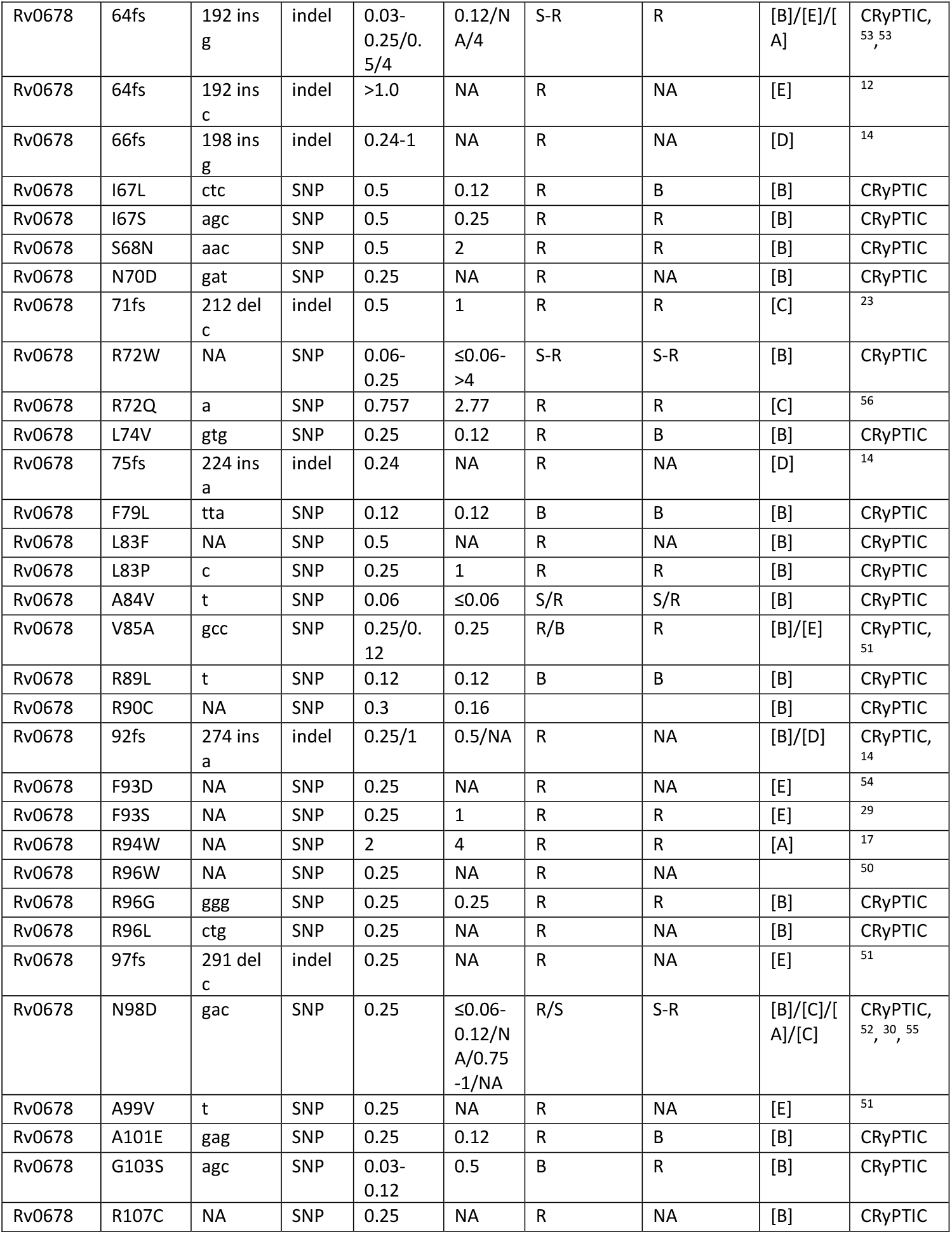
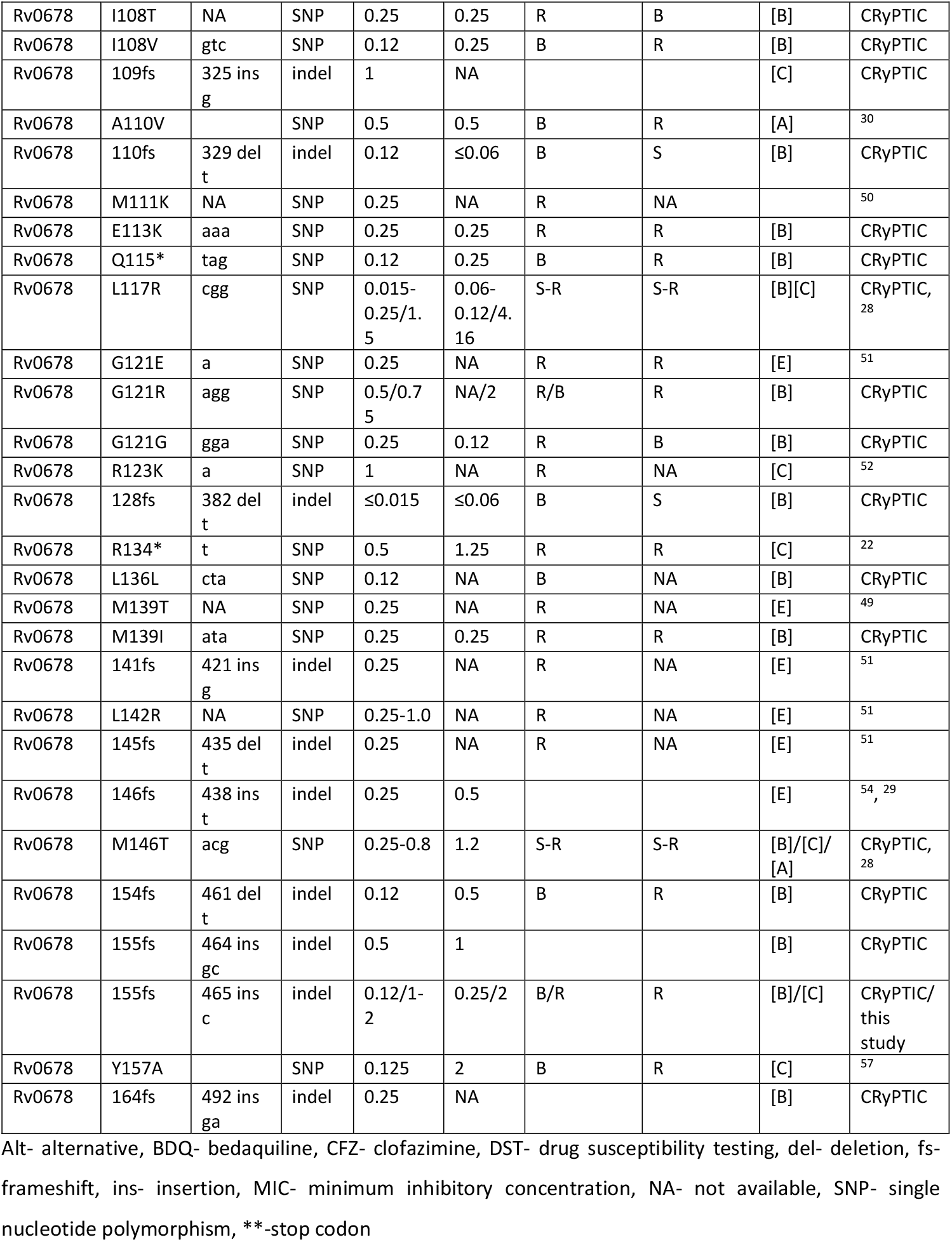
*Rv0678* mutations detected in patients (published literature and CRyPTIC collection). Reference catalog of *Rv0678* mutations which have been described in previously published patient isolates or collected by the Comprehensive Resistance Prediction for Tuberculosis an International Consortium (CRyPTIC) partners. Additionally, alternative (Alt.) allele information included, describes the variants at the base pair position. Variants included in list if presented a borderline or resistant phenotype to at least bedaquiline (BDQ)MIC determined by different drug susceptibility testing (DST) techniques depending on the study, abbreviated as: [A] BD-MGIT™, [B] UKMYC5/UKMYC6 plates, [C] Alamar blue/resazurin assay, [D] 7H10 plates, [E] 7H11 plates Phenotype interpretation was based off of: [A] 1mg/L BDQ/CFZ^45^; [B] 0.12-0.25mg/L BDQ/CFZ^46^, [C] defined by study authors, [D] 0.5mg/L BDQ^47^/CFZ^48^, [E] 0.25 mg/L BDQ^45^-CFZ determined by study authors

| Gene | Substitution | Alt Allele | Type | BDQ MIC (mg/L) | CFZ MIC (mg/L) | Phenotype interpretation on BDQ- R, B, S | Phenotype interpretation on CFZ- R, B, S | DST Method | Author |
| --- | --- | --- | --- | --- | --- | --- | --- | --- | --- |
| Rv0678 | V1A | c | SNP | 6.4/0.5 | >4/NA | R/B | R | [A] | <sup>24, 25</sup> |
| Rv0678 | S2I | t | SNP | 0.73 | 4 | R | R | [C] | <sup>28</sup> |
| Rv0678 | N4T | acc | SNP | 0.06-0.12 | 0.12 | S/B | B | [B] | CRyPTIC |
| Rv0678 | 5fs | 15 del c | indel | 0.5 | 0.25 | R | R | [B] | CRyPTIC |
| Rv0678 | 6fs | 18 del gg | indel | 0.5 | NA | R | NA | [E] | <sup>49</sup> |
| Rv0678 | 6fs | 16 del gg | indel | 0.5 | NA | R | NA |  | <sup>50</sup> |
| Rv0678 | 10fs | 28 del t | indel | 0.12 | ≤0.06 | B | S | [B] | CRyPTIC |
| Rv0678 | 10fs | 29 del t | indel | 0.25 | 0.12 | R | B | [B] | CRyPTIC |
| Rv0678 | M10I | atc | SNP | 0.12 | NA | B | NA | [B] | CRyPTIC |
| Rv0678 | 11fs | 32 del g | indel | 0.25 | NA | R | NA | [E] | <sup>51</sup> |
| Rv0678 | G11G | t | SNP | 0.5-2.0 | NA | R | NA | [C] | <sup>52</sup> |
| Rv0678 | P14S | tcc | SNP | 0.25 | 0.5 | R | R | [B] | CRyPTIC |
| Rv0678 | P14L | ctc | SNP | 0.12 | ≤0.06 | B | S | [B] | CRyPTIC |
| Rv0678 | D15G | NA | SNP | 0.25 | NA | R | NA | [B] | CRyPTIC |
| Rv0678 | F19S | tcc | SNP | 0.5 | NA | R | NA | [B] | CRyPTIC |
| Rv0678 | V20G | NA | SNP | 0.25/2/0.25/0.25 | NA/2/NA/4 | R/B | R | [E]/[A]/[A]/[E] | <sup>53, 53, 54, 29</sup> |
| Rv0678 | V20A | gcc | SNP | 0.12 | 0.12 | B | B | [B] | CRyPTIC |
| Rv0678 | E21D | c | SNP | 0.5-1.0 | NA | R | NA | [C] | <sup>52</sup> |
| Rv0678 | S29S | tct | SNP | 0.12 | 0.12 | B | B | [B] | CRyPTIC |
| Rv0678 | S31R | c | SNP | 0.5 | NA | R | NA | [C] | <sup>52</sup> |
| Rv0678 | L32S | tcg | SNP | 0.12 | 0.12 | B | B | [B] | CRyPTIC |
| Rv0678 | T33A | g | SNP | 0.5 | 2 | R | R | [C] | <sup>23</sup> |
| Rv0678 | T33S | g | SNP | 2 | NA | R | NA | [C] | <sup>52</sup> |
| Rv0678 | A36V | t | SNP | 0.5 | 4 | R | R | [C] | <sup>23</sup> |
| Rv0678 | 36fs | 107 ins<br>g | indel | 0.5 | 0.25 | R | R | [B] | CRyPTIC |
| Rv0678 | L40M | atg | SNP | 0.5 | NA | R | NA | [B] | CRyPTIC |
| Rv0678 | L40F | ttc | SNP | 0.5 | NA | R | R | [B] | CRyPTIC |
| Rv0678 | W42R | cgg | SNP | 0.25/0.12 | 0.25/2 | R/B | R | [B]/[D] | CRyPTIC,<br><sup>14</sup> |
| Rv0678 | 46fs | 136 ins<br>g | indel | 1 | NA | R | NA | [E] | <sup>54</sup> |
| Rv0678 | 46fs | 137 ins<br>g | indel | 0.25 | 1 | R | R | [B] | CRyPTIC |
| Rv0678 | 46fs | 138 ins<br>g | indel | 0.25/0.25/2/>1/1 | 0.25/NA/4/NA/NA | R | R | [B]/[E]/[A]/[E]/[E] | CRyPTIC,<br><sup>53 53 12 54</sup> |
| Rv0678 | 46fs | 138 ins<br>ga | indel | >1.0 | NA | R | NA | [E] | [15] |
| Rv0678 | C46R | cgt | SNP | 0.25 | NA | R | NA | [B] | CRyPTIC |
| Rv0678 | 47fs | 139 ins<br>g | indel | 0.25 | NA | R | NA | [E] | <sup>51</sup> |
| Rv0678 | 47fs | 140 ins<br>g | indel | 0.25 | NA | R | NA | [E] | <sup>49</sup> |
| Rv0678 | 47fs | 141 ins<br>c | indel | 0.06-0.25/0.5/4/0.5 | 0.06-1/NA/4/NA | S-R | S-R | [B]/[E]/[A]/[E] | CRyPTIC,<br><sup>53 53 29</sup> |
| Rv0678 | 48fs | 144 ins<br>c | indel | 0.25 | NA | R | NA | [E] | <sup>51</sup> |
| Rv0678 | P48L | NA | SNP | 0.25-0.5 | NA | R | B | [B] | CRyPTIC |
| Rv0678 | E49fs | NA | indel | 0.03-0.25 | 0.25-2 | R | R | [E] | <sup>29</sup> |
| Rv0678 | R50Q | NA | SNP | 0.25 | NA | R | NA | [B] | CRyPTIC |
| Rv0678 | Q51K | NA | SNP | 0.25 | 1 | R | R | [B] | CRyPTIC |
| Rv0678 | Q51R | NA | SNP | 0.3 | 1 | R | R | [B] | CRyPTIC |
| Rv0678 | S52F | NA | SNP | 0.48 | NA | R | NA | [D] | <sup>14</sup> |
| Rv0678 | S53L | NA | SNP | 0.25-0.4 | 2 | R | R | [C] | <sup>28</sup> |
| Rv0678 | S53P | NA | SNP | 0.5 | 2-4 | R | R | [C] | <sup>28</sup> |
| Rv0678 | 58fs | 172 ins<br>IS6110 | indel | 0.5 | 1 | R | R | [C] | <sup>23</sup> |
| Rv0678 | A59V | NA | SNP | 0.48 | NA | R | NA | [D] | <sup>14</sup> |
| Rv0678 | A62V | NA | SNP | 0.48 | NA | R | NA | [D] | <sup>14</sup> |
| Rv0678 | A62T | acc | SNP | 0.12 | 0.5 | B | R | [B] | CRyPTIC |
| Rv0678 | S63R | a | SNP | 0.5 | 1.25 | R | R | [C] | <sup>22</sup> |
| Rv0678 | S63N | a | SNP | 0.25 | NA | R | NA | [C] | <sup>55</sup> |
| Rv0678 | 64fs | 192 ins<br>g | indel | 0.03-0.25/0.<br>5/4 | 0.12/N<br>A/4 | S-R | R | [B]/[E]/[<br>A] | CRyPTIC,<br>53 53<br>, |
| Rv0678 | 64fs | 192 ins<br>c | indel | >1.0 | NA | R | NA | [E] | 12 |
| Rv0678 | 66fs | 198 ins<br>g | indel | 0.24-1 | NA | R | NA | [D] | 14 |
| Rv0678 | I67L | ctc | SNP | 0.5 | 0.12 | R | B | [B] | CRyPTIC |
| Rv0678 | I67S | agc | SNP | 0.5 | 0.25 | R | R | [B] | CRyPTIC |
| Rv0678 | S68N | aac | SNP | 0.5 | 2 | R | R | [B] | CRyPTIC |
| Rv0678 | N70D | gat | SNP | 0.25 | NA | R | NA | [B] | CRyPTIC |
| Rv0678 | 71fs | 212 del<br>c | indel | 0.5 | 1 | R | R | [C] | 23 |
| Rv0678 | R72W | NA | SNP | 0.06-0.25 | ≤0.06-<br>>4 | S-R | S-R | [B] | CRyPTIC |
| Rv0678 | R72Q | a | SNP | 0.757 | 2.77 | R | R | [C] | 56 |
| Rv0678 | L74V | gtg | SNP | 0.25 | 0.12 | R | B | [B] | CRyPTIC |
| Rv0678 | 75fs | 224 ins<br>a | indel | 0.24 | NA | R | NA | [D] | 14 |
| Rv0678 | F79L | tta | SNP | 0.12 | 0.12 | B | B | [B] | CRyPTIC |
| Rv0678 | L83F | NA | SNP | 0.5 | NA | R | NA | [B] | CRyPTIC |
| Rv0678 | L83P | c | SNP | 0.25 | 1 | R | R | [B] | CRyPTIC |
| Rv0678 | A84V | t | SNP | 0.06 | ≤0.06 | S/R | S/R | [B] | CRyPTIC |
| Rv0678 | V85A | gcc | SNP | 0.25/0.<br>12 | 0.25 | R/B | R | [B]/[E] | CRyPTIC,<br>51 |
| Rv0678 | R89L | t | SNP | 0.12 | 0.12 | B | B | [B] | CRyPTIC |
| Rv0678 | R90C | NA | SNP | 0.3 | 0.16 |  |  | [B] | CRyPTIC |
| Rv0678 | 92fs | 274 ins<br>a | indel | 0.25/1 | 0.5/NA | R | NA | [B]/[D] | CRyPTIC,<br>14 |
| Rv0678 | F93D | NA | SNP | 0.25 | NA | R | NA | [E] | 54 |
| Rv0678 | F93S | NA | SNP | 0.25 | 1 | R | R | [E] | 29 |
| Rv0678 | R94W | NA | SNP | 2 | 4 | R | R | [A] | 17 |
| Rv0678 | R96W | NA | SNP | 0.25 | NA | R | NA |  | 50 |
| Rv0678 | R96G | ggg | SNP | 0.25 | 0.25 | R | R | [B] | CRyPTIC |
| Rv0678 | R96L | ctg | SNP | 0.25 | NA | R | NA | [B] | CRyPTIC |
| Rv0678 | 97fs | 291 del<br>c | indel | 0.25 | NA | R | NA | [E] | 51 |
| Rv0678 | N98D | gac | SNP | 0.25 | ≤0.06-0.12/N<br>A/0.75<br>-1/NA | R/S | S-R | [B]/[C]/[<br>A]/[C] | CRyPTIC,<br>52, 30, 55 |
| Rv0678 | A99V | t | SNP | 0.25 | NA | R | NA | [E] | 51 |
| Rv0678 | A101E | gag | SNP | 0.25 | 0.12 | R | B | [B] | CRyPTIC |
| Rv0678 | G103S | agc | SNP | 0.03-0.12 | 0.5 | B | R | [B] | CRyPTIC |
| Rv0678 | R107C | NA | SNP | 0.25 | NA | R | NA | [B] | CRyPTIC |
| Rv0678 | I108T | NA | SNP | 0.25 | 0.25 | R | B | [B] | CRyPTIC |
| Rv0678 | I108V | gtc | SNP | 0.12 | 0.25 | B | R | [B] | CRyPTIC |
| Rv0678 | 109fs | 325 ins<br>g | indel | 1 | NA |  |  | [C] | CRyPTIC |
| Rv0678 | A110V |  | SNP | 0.5 | 0.5 | B | R | [A] | <sup>30</sup> |
| Rv0678 | 110fs | 329 del<br>t | indel | 0.12 | ≤0.06 | B | S | [B] | CRyPTIC |
| Rv0678 | M111K | NA | SNP | 0.25 | NA | R | NA |  | <sup>50</sup> |
| Rv0678 | E113K | aaa | SNP | 0.25 | 0.25 | R | R | [B] | CRyPTIC |
| Rv0678 | Q115* | tag | SNP | 0.12 | 0.25 | B | R | [B] | CRyPTIC |
| Rv0678 | L117R | cgg | SNP | 0.015-0.25/1.<br>5 | 0.06-0.12/4.<br>16 | S-R | S-R | [B][C] | CRyPTIC,<br><sup>28</sup> |
| Rv0678 | G121E | a | SNP | 0.25 | NA | R | R | [E] | <sup>51</sup> |
| Rv0678 | G121R | agg | SNP | 0.5/0.7<br>5 | NA/2 | R/B | R | [B] | CRyPTIC |
| Rv0678 | G121G | gga | SNP | 0.25 | 0.12 | R | B | [B] | CRyPTIC |
| Rv0678 | R123K | a | SNP | 1 | NA | R | NA | [C] | <sup>52</sup> |
| Rv0678 | 128fs | 382 del<br>t | indel | ≤0.015 | ≤0.06 | B | S | [B] | CRyPTIC |
| Rv0678 | R134* | t | SNP | 0.5 | 1.25 | R | R | [C] | <sup>22</sup> |
| Rv0678 | L136L | cta | SNP | 0.12 | NA | B | NA | [B] | CRyPTIC |
| Rv0678 | M139T | NA | SNP | 0.25 | NA | R | NA | [E] | <sup>49</sup> |
| Rv0678 | M139I | ata | SNP | 0.25 | 0.25 | R | R | [B] | CRyPTIC |
| Rv0678 | 141fs | 421 ins<br>g | indel | 0.25 | NA | R | NA | [E] | <sup>51</sup> |
| Rv0678 | L142R | NA | SNP | 0.25-1.0 | NA | R | NA | [E] | <sup>51</sup> |
| Rv0678 | 145fs | 435 del<br>t | indel | 0.25 | NA | R | NA | [E] | <sup>51</sup> |
| Rv0678 | 146fs | 438 ins<br>t | indel | 0.25 | 0.5 |  |  | [E] | <sup>54</sup>, <sup>29</sup> |
| Rv0678 | M146T | acg | SNP | 0.25-0.8 | 1.2 | S-R | S-R | [B]/[C]/<br>[A] | CRyPTIC,<br><sup>28</sup> |
| Rv0678 | 154fs | 461 del<br>t | indel | 0.12 | 0.5 | B | R | [B] | CRyPTIC |
| Rv0678 | 155fs | 464 ins<br>gc | indel | 0.5 | 1 |  |  | [B] | CRyPTIC |
| Rv0678 | 155fs | 465 ins<br>c | indel | 0.12/1-2 | 0.25/2 | B/R | R | [B]/[C] | CRyPTIC/<br>this<br>study |
| Rv0678 | Y157A |  | SNP | 0.125 | 2 | B | R | [C] | <sup>57</sup> |
| Rv0678 | 164fs | 492 ins<br>ga | indel | 0.25 | NA |  |  | [B] | CRyPTIC |

Alt- alternative, BDQ- bedaquiline, CFZ- clofazimine, DST- drug susceptibility testing, del- deletion, fs-frameshift, ins- insertion, MIC- minimum inhibitory concentration, NA- not available, SNP- single nucleotide polymorphism, \*-stop codon

Wild-type isolates presented an MIC95 of 0.12 mg/L BDQ/CFZ in the UKMYC5/6 plates^58^. Therefore, the patient isolates in the study were catalogued as >0.12 mg/L BDQ/CFZ resistant, <0.12 susceptible, and =0.12 borderline. The CRyPTIC dataset contained 178 patient isolates with a single *Rv0678* variant, 52 of which were classified as BDQ resistant (27 cross-resistant with CFZ), 32 with borderline BDQ resistance, and 94 BDQ susceptible (Table S4). To better explain this high number of BDQ susceptible strains with *Rv0678* mutations we looked for additional mutations in the efflux pump genes *mmpL5* and *mmpS5*, as it has been postulated that variants in these genes may reduce or abrogate acquired resistance^59^. However, we found that 64/94 (68%) of susceptible isolates harboring *Rv0678* mutations had WT *mmpL5/mmpS5* genes (Table S4). Likely functional/inactivating mutations in *mmpL5* (19/19 isolates with frameshifts and 3/5 with missense) were associated with susceptibility while synonymous mutations appeared uncorrelated with susceptibility (7/18 isolates susceptible).

Plotting all 147 resistance and borderline resistance-associated *Rv0678* mutations on the gene sequence (Figure 1), reveals that they occur throughout the entire length of the gene with no single hot spot region. However, there are several regions where mutations are more likely to occur, i.e. codons 46-53 and 62-68 (Figure 3B).

Next, we searched the CRyPTIC dataset for resistance conferring mutations (BDQ MIC >0.12 mg/L) in *atpE, pepQ* (*Rv2535c*) and *Rv1979c*. After filtering out phylogenetic SNPs and strains with preexisting *Rv0678* mutations, we found 35 isolates with an MIC >0.12 mg/L for BDQ harboring *Rv1979c* mutations and only 3 isolates with mutations in *atpE*^60^. There were no BDQ resistant isolates with *pepQ* mutations, but 19 strains had borderline BDQ resistance with an MIC of 0.12 mg/L. In total, we defined four *Rv1979c*-mediated resistant mutations, and 7 possible borderline-resistance conferring mutations in *pepQ* (*Rv2535c*).

Finally, after an in-depth literature search including *in vitro*, *in vivo*, and patient derived strains, we complied a comprehensive catalogue of variants in BDQ resistance associated genes (Table S5). Due to insufficient data for BDQ and CFZ critical concentration determination for many phenotypic assays, and the high overlap of MIC distributions between WT and mutant clones, resistance interpretation of mutations was based on the ascribed phenotype in each publication or by WHO critical concentration when available.

### Structural variation in Rv0678

For better mechanistic understanding of how *Rv0678* mutations result in BDQ/CFZ resistance, we mapped missense mutations from our catalogue to the previously determined Rv0678 crystal structure (107 mutations mapped to resolved structure positions PDB ID 4NB5, Table S5)^20^. As mutations in different parts of the protein structure are likely to have different effects, we investigated four different mechanisms for conferring resistance: reducing folded protein/protein stability, impairing DNA binding, altering protein dimerization, and reducing affinity for the fatty acid ligand of Rv0678.

The numerous introductions of premature stop codons and frameshift mutations reported here and elsewhere suggest loss of functional Rv0678 protein is a common mechanism of resistance^22,41,51^. Consistent with this, if we compare the missense *Rv0678* mutations in resistant isolates with those in susceptible isolates, we find the former are more commonly predicted to destabilize the Rv0678 wHTH domain required for DNA binding (destabilization defined as ΔΔG <= −1 kcal/mol as measured by mCSM protein stability, 34/38 R, 5/10 S, OR 7.96, Fisher exact p=0.012), suggesting that disruption of protein stability is also a resistance mechanism. These findings are also compatible with the observation that missense mutations from resistant isolates are predicted to have a more deleterious effect by SNAP2 score compared with those from susceptible isolates (Wilcoxon p=0.018)^61^.

To understand how mutations may impact DNA binding, we structurally aligned the wHTH DNA binding domain of Rv0678 to several other Mtbc marR-family transcriptional regulators whose DNA-bound structures have been determined (Figure 4B, S3; average RMSD 0.81 Å)^62^. We found that mutations clustering in regions 62-68 and 88-92 correlate with the recognition α-helix and conserved DxR motif respectively, which directly contact the DNA and have previously been shown to critical for the binding affinity of other marR-family regulators (Figure 4B)^62,63^. However, further experimental verification of these mutations’ effects on DNA binding is required.

**Figure 4:**
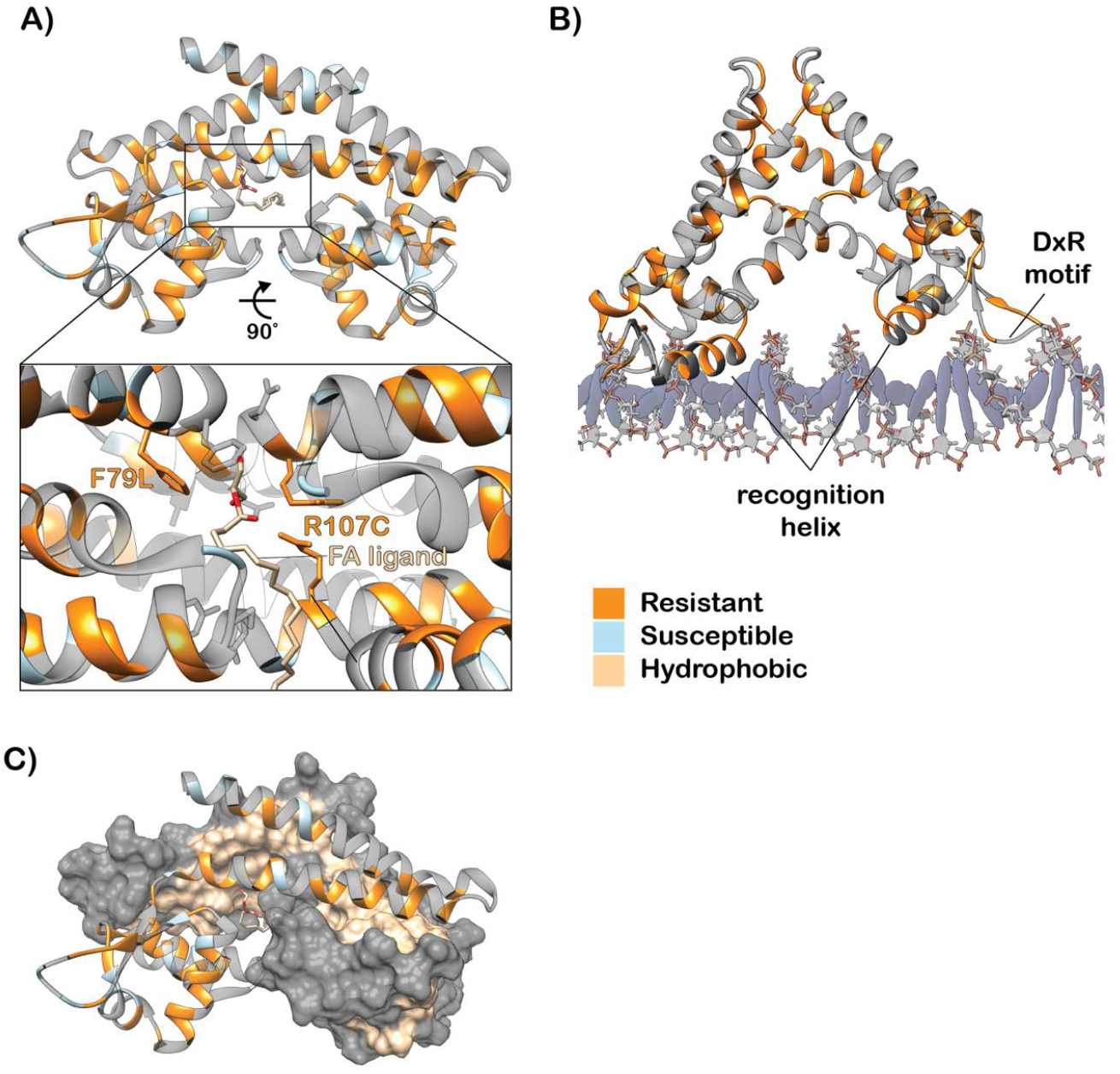
Structural mechanisms of BDQ resistance. (A) Rv0678 dimer with resistant (orange) and susceptible (blue) mutations shown. Pullout highlights the resistant mutations in the ligand binding pocket. (B) Rv0678 dimer modeled onto DNA. Conserved MarR-family DNA binding elements are noted. (C) Resistant mutations occur across the hydrophobic dimerization domain.

We also investigated the possibility that mutations alter the protein-protein interactions of the Rv0678 dimer. Mutations associated with resistance are enriched at the dimer interface (27/29 interface mutations R), suggesting they may impact dimer formation and/or stability (Figure 4C). However, computational prediction of interface mutations’ effects on non-DNA bound dimer stability revealed no significant difference in destabilization as defined by mCSM PPI ΔΔG <= −1 kcal/mol (16/27 R, 1/2 S). Interpretation of the DNA-bound dimer interface was not feasible due to the significant conformational rearrangement of the dimer alpha-helices upon DNA binding. Finally, all isolates where an amino acid that contacts the fatty acid ligand 2-stearoylglycerol was mutated were resistant, suggesting that disrupting the binding of the ligand confers resistance, possibly through perturbing non-DNA bound homodimer formation (Figure 4A).

### Molecular dynamics simulations

Rv0678 must also undergo a conformational rearrangement to bind DNA which is not captured by the static structures in the previous section. Therefore, we employed molecular dynamics simulations to understand how particular mutations in the “hinge” region (connecting the DNA binding and dimerization domains) might disrupt this motion and lead to resistance. To do this, we performed 100ns simulations on the WT Rv0678 homodimer as well as Rv0678 homodimers containing either L40F and A101E mutations (which are associated with resistance but do not clearly interact with the DNA or disrupt protein folding) or the L40V mutation (consistently phenotypically susceptible). These simulations revealed that mutations from resistant isolates (L40F and A101E) resulted in changes in the conformational dynamics and pocket crosstalk of the individual Rv0678 monomers distinct from WT or the L40V mutation (Figure 5). In particular, A101E formed a new stable salt-bridge interaction in the “hinge” domain, which could disrupt the conformational flexibility required by this region to transition between the DNA- and ligand-bound homodimer states (Figure S4).

**Figure 5:**
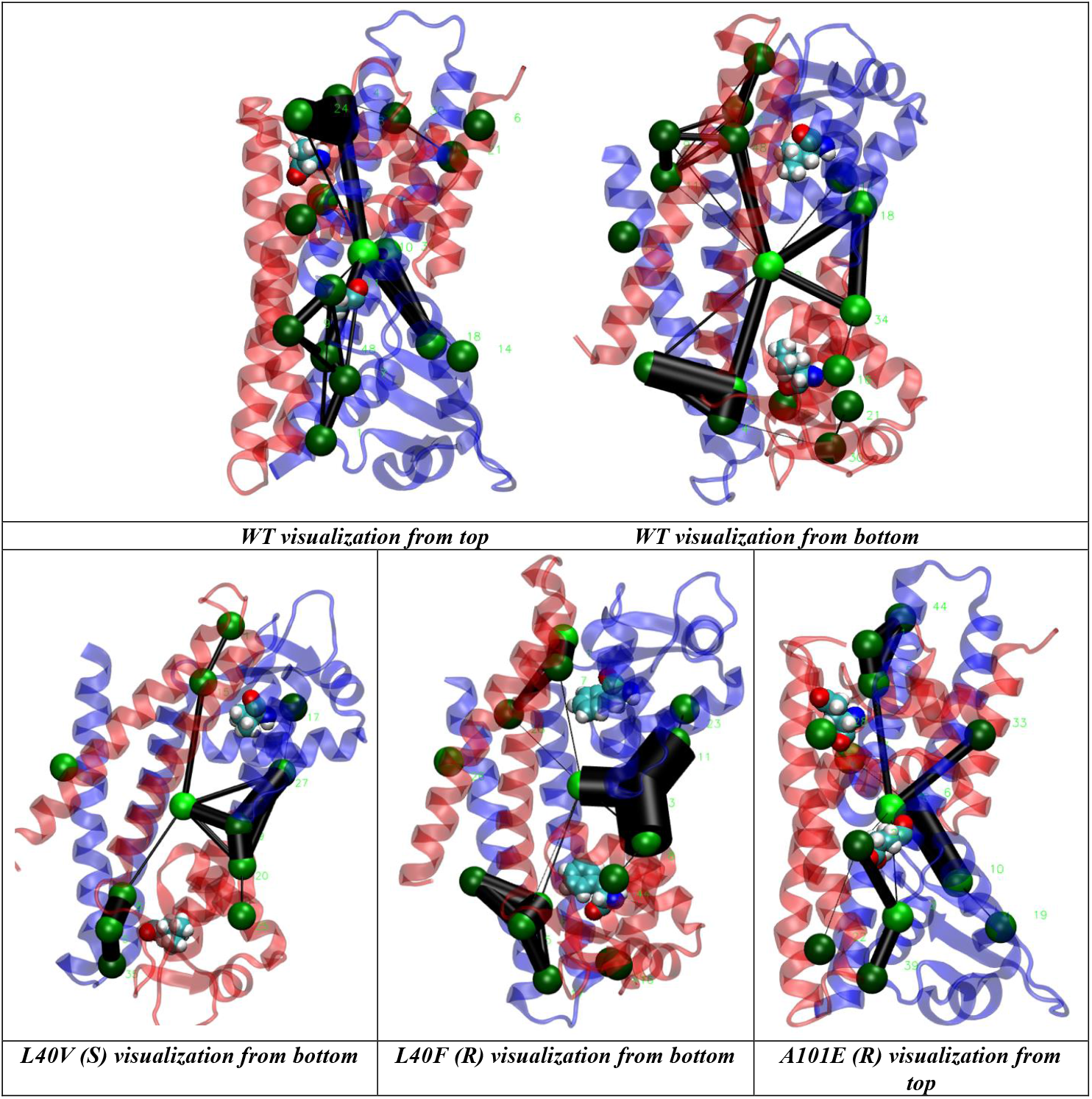
Networks of the most persistent pockets found in the wild-type (WT) and mutated Rv0678. The pocket crosstalk analysis on simulated systems identifies allosteric signaling. All systems shared a large common pocket located at the interface of the two monomers (Figure S5). This large pocket communicates with other smaller pockets at different zones of the homodimer by the network edges (black lines). Comparative analysis showed that the resistance-conferring mutations, L40F and A101E displayed different network edges during the 100ns of simulations with respect to the WT. R-resistant, S-susceptible

## Discussion

In this study, we identify new bedaquiline and clofazimine resistance determinants by combining *in vitro* evolution experiments with WGS and MIC data, including the analysis of 14,151 clinical Mtbc strains provided by the CRyPTIC consortium (http://www.crypticproject.org/). The results have allowed us to define a comprehensive set of genomic variants associated with resistance to BDQ (& CFZ), which are central to treating MDR, pre-XDR and XDR-TB^8^. This project also demonstrates the potential of *in vitro* evolution experiments employing selection under sub-lethal drug concentrations to generate clones with a wide spectrum of mutations, including clinically relevant mutations. Collectively, this work helps extend the use of genomics in providing personalized medicine to patients with MDR, pre-XDR, and XDR-TB, through bolstering genotypic catalogues with phenotypically verified variant lists.

Although BDQ/CFZ are rolled out worldwide as core drugs in treatment regimens for MDR-TB patients^8^, antibiotic susceptibility testing for these compounds is rarely available and patients are often treated empirically. Indeed, recent studies report treatment failure and/or poor outcome of MDR-TB patients as mutations in *Rv0678* emerge that are associated with BDQ/CFZ resistance^12,15,17,26,54^. Additionally, misdiagnosis can promote the selection and subsequent transmission of specific MDR Mtbc strains. For example, it was recently reported in Eswatini and South Africa, that patients infected with an MDR outbreak strain harboring the RMP mutation I491F, which is not detectable by conventional phenotypic or genotypic testing, probably continued to receive RMP although it was ineffective^30^. These strains now represent more than 60% of MDR strains in Eswatini and additionally more than half of these developed enhanced BDQ/CFZ MICs via acquired *Rv0678* mutations^1,29,30,64^ This scenario underlines the urgent need for approaches to rapidly detect resistance, including to BDQ/CFZ, with baseline diagnostics and during therapy (e.g. via WGS or targeted genome sequencing) to avoid treatment failure, further resistance development and ongoing transmission of specific MDR strains.

Genome sequencing techniques have been demonstrated to accurately identify genomic variants in all target regions involved in resistance development in a single analysis^3,65^. However, several major challenges exist—including linking genotype and phenotype to distinguish resistance-mediating from benign mutations and interpreting how multiple mutations interact when the resistance phenotype is polygenic, as is the case with BDQ and CFZ^60^. This work catalogues the phenotype of 260 unique genomic variants across six genes. Importantly, not all variation leads to resistance, as 38 of these mutations in *Rv0678* are not associated with resistance to BDQ/CFZ (Table S5), which facilitates the prediction of BDQ/CFZ susceptibility. Furthermore, 30/94 BDQ susceptible isolates with *Rv0678* mutations harbored an additional mutation in the *mmpL5* or *mmpS5* efflux pump genes, which may abrogate the efflux pump mechanism and reconstitute the susceptible phenotype^59^. Here, we provide preliminary evidence that isolates with inactivating (frameshift, nonsense, or missense variants) *mmpL5* mutations can override *Rv0678* resistance-mediating mutations, resulting in hyper-susceptibility to BDQ. Further work to understand the frequency, distribution and origin of these inactivating mutations is necessary to understand their clinical relevance worldwide.

While genome sequencing of clinical Mtbc strains has been deployed in several countries, its implementation is constrained not just by technical and data analysis considerations, but also because working directly from clinical samples (e.g. sputum) is difficult^3^. We recently demonstrated that targeted genome sequencing using the commercially available Deeplex^®^-MycTB has a high sensitivity and specificity for first- and second-line drug resistance prediction, including BDQ/CFZ resistance mutations that have been found in patients^64,66,67^. In combination with our comprehensive set of mutations that confer resistance to BDQ/CFZ, Deeplex^®^-MycTB and other targeted sequencing approaches allow for rapid detection of BDQ/CFZ resistance from clinical samples in few days using small portable genome sequencers such as the MinION^68^.

Importantly, *in vitro* laboratory evolution experiments identified 50 resistance-conferring mutations that have not yet appeared in clinical strains, thus allowing us to characterize and identify a significant proportion of resistance-conferring mutations before they have even occurred in a patient. These results highlight the value of *in vitro* evolution experiments as a complementary method for prospective identification of resistance, as compared to traditional surveillance of patient isolates. Furthermore, this method provides a large strain collection of *de novo* resistant clones from a comparable wild-type susceptible ancestor. Finally, by using PacBio sequencing to establish the full genome sequence of mutant clones without any mutations in the relevant targets, we were able to identify a large genomic inversion that disrupts *Rv0678*, which should be considered as a novel BDQ/CFZ resistance mechanism.

Another method for prospectively identifying mutations that confer resistance is structure-based modeling, which has recently been combined with machine learning methods to design algorithms for predicting resistance to rifampicin, isoniazid, and pyrazinamide^69–73^. These approaches rely on the fact that resistance-conferring mutations are often located in drug binding pockets or active sites and have distinct biophysical consequences as compared to their susceptible counterparts^74^. This study identifies several resistance mechanisms in *Rv0678* with quantifiable structural effects (protein stability, dimer interactions, SNAP2 scores, and interaction with the DNA), suggesting that a structure-based machine learning approach could also be successful for predicting BDQ/CFZ resistance.

As we have shown, any approach to predict BDQ resistance must consider not only mutations present in *Rv0678*, but also potential resistance/susceptibility determinants in other putative resistance-conferring genes e.g. *atpE, pepQ, Rv1979c*, and *mmpL/S5*. It has been attempted to predict BDQ resistance from *atpE* mutations^75^, so one feasible option would be a meta-approach that flags if any individual gene-level algorithms predict resistance/susceptibility. In addition, CRyPTIC and other studies are producing large MIC datasets, which will enable much more nuanced modeling of resistance, which is particularly important for drugs with efflux-mediated resistance that often exhibit small sub-threshold changes in MIC. Although structure-based modeling is helpful in predicting resistance, these structural hypotheses must be confirmed experimentally for final validation.

As indicated by the overlap of *in vitro* selected mutations with 11 resistance conferring mutations found in patients, sub-MIC drug concentration can select for clinically relevant resistance variants. This modeling might also better imitate the intra-patient situation, since the complex pathology of TB disease prevents certain drugs (i.e. bactericidal and bacteriostatic drugs), from reaching all bacteria at a therapeutic level (for example structures like pulmonary cavities and granuloma lesions)^33,35,36,77^. BDQ and CFZ in particular do not penetrate the central caseous lesion (or necrotic center) of the granuloma effectively^34^. Although this study provides a large collection of *in vitro* selected Rv0678-mediated resistant clones, the efflux pump modulation by these various mutations was not experimentally investigated and should be explored in the future.

In addition to clear-cut resistance-associated mutations, additional “secondary” mutations were coselected with resistance-associated variants *in vitro*. These secondary mutations affected several genes, mainly associated with cellular processes such as cell wall synthesis, metabolic processes, and protein regulation; with mutations also in genes of unknown function (Table S3). It is possible that these mutations bring about an additional phenotypic affect, or they may be compensatory mutations^5,75^. However, until further experiments are conducted, such as competitive fitness assays (or even transcriptomic analysis), it cannot be ruled out that these are merely hitchhiking mutations or culture-selected variants.

In this study, we focused on structural mechanisms of *Rv0678* mediated resistance, as these mutations are the most clinically relevant and correlated with BDQ treatment failures. Other genes which have been linked to BDQ or CFZ resistance, such as *atpE, pepQ* and *Rv1979c*, but their clinical relevance has yet to be established^10,29,49,53^. In the CRyPTIC dataset we identified four *Rv1979c* variants which presented a resistant phenotype to BDQ with variable CFZ cross resistance, and seven mutations in *pepQ* garnering borderline BDQ resistance and variable CFZ resistance. Finally, mutations in *atpE* (coding for the target of BDQ) were rare among the *in vitro* selection experiments and the CRyPTIC dataset, suggesting that *atpE* mutations might have a high fitness cost.

In conclusion, our work advances our understanding of how resistance can arise to BDQ and CFZ. This information provides an updated variant catalogue, comprising of mutations and structural variations associated with BDQ/CFZ resistance as well as benign variants and mutations implicated with hyper-susceptibility. Employing this information in DNA sequencing-based diagnostic approaches will allow for the first time a rational design of BDQ and CFZ-containing MDR-TB therapies.

## Materials and Methods

### *In vitro* evolution experiments

*In vitro* experiments were carried out with H37Rv strain (ATCC 27294). Bedaquiline was purchased from Janssen-Cilag GmbH and clofazimine was ordered from Sigma (C8895-1G) both were reconstituted from powder in DMSO and stored at −20°C.

Evolutionary experiments were conducted with the *Mycobacterium tuberculosis* complex (Mtbc) lab strain H37Rv. The bacteria were cultured from frozen stocks and (pre)cultured at 37° in 7H9 medium, supplemented with 0.2% glycerol, 10% oleic albumin dextrose catalase (OADC), and 0.05% Tween^80^(termed culture medium). At exponential growth phase (between 0.3 and 0.6 optical density (OD) of 600nm), bacteria were transferred to new culture medium with a final OD of 0.05 in 50 mL, additionally supplemented with either bedaquiline (BDQ) or clofazimine (CFZ). Four concentrations below the minimum inhibitory concentration (MIC) were included for each drug at 1:2, 1:4, 1:8, and 1:16 below the strain MIC (and an antibiotic-free control). The MIC of the susceptible WT ancestor was determined to be 0.12 mg/L BDQ and 0.5 mg/L CFZ in this culturing system.

These experiments were carried out over 20 days with five culture passages total. Culture passages were conducted every four days with a final culture OD of 0.05 transferred. After passages 1, 3, and 5; bacteria were plated on selective agar plates of 7H11, supplemented with 0.5% glycerol, 10% OADC and either 0.12 (1:2 MIC) or 0.25 (MIC) mg/L of BDQ, or 0.25 (1:2 MIC) mg/L CFZ. After 21-28 days of growth on selective agar plates, single colonies were transferred to culture medium, grown for 2-4 weeks, then two 1 mL aliquots were frozen at −80, and samples were taken for whole genome sequencing (WGS). All remaining colonies (collected by experimental condition) were collected for WGS, i.e. total population sequencing.

### Whole genome sequencing

Isolates from *in vitro* evolutionary experiments underwent DNA isolation by CTAB method^79^. Paired-end DNA library preparations and sequencing was performed with Illumina technology (Nextera-XT and NextSeq500) according to the manufacturer’s instructions and with a minimum average genome coverage of 50x. Fastq files (raw sequencing data) were mapped to the *M. tuberculosis* H37Rv reference genome (GenBank ID: NC_000962.3) using the MTBseq pipeline^80^.

Briefly, for the analysis of single mutants we considered single nucleotide polymorphisms (SNPs) which were covered by a minimum of four reads in both forward and reverse orientation, four reads calling the allele with at least a phred score of 20, and an allele frequency of 75%. For the detection of insertions and deletion (InDels) we allowed a frequency over 50%.

Deep sequencing was conducted for population diversity analysis with an average genome wide coverage of 200 – 420; SNPs and InDels were called with at least one forward and one reverse read and a phred score >20 for at least 2 reads.

Three isolates underwent sequencing using the PacBio Sequel II System (Pacific Biosciences). Libraries were prepared with the SMRTbell^®^ Express Template Prep Kit 2.0 according to the manufacturer’s instructions. Barcoded overhang adapters for multiplexing were ordered at IDT (Integrated DNA Technologies). During demultiplexing barcodes were filtered for a minimum quality of 50t, which yielded long read sequencing data of an average of 4.4 GB and mean subread length of 9 KB. Long read sequences were *de novo* assembled using PacBio SMRT^®^ Link software version 9 and the “Microbial Assembly” workflow with a set genome size of 4.5 GB with default parameters.

### Screening of Rv0678 mutations in clinical samples via the CRyPTIC strain collection

Patient derived Mtbc isolates were collected by CRyPTIC partners throughout 27 countries and analyzed in 14 different laboratories. Strains were selected from the CRyPTIC database for this study if they had matched genotype and phenotype data, a high quality BDQ phenotype, and a mutation in BDQ resistance associated gene (either *atpE, Rv1979c, pepQ, mmpL/S5*, or *Rv0678*). This led to the curation of 179 strains in total.

The full analysis pipeline for CRyPTIC is documented in [paper in preparation], but we outline the key steps here. Sequencing reads were deposited at the European Bioinformatics Institute and run through a bespoke bioinformatics pipeline (publicly available here: https://github.com/iqbal-lab-org/clockwork). In short, reads were filtered against human and other microbial species reads before being mapped to the H37Rv reference genome. Two parallel variant callers (SamTools and Cortex) were used^81,82^, one of which makes high sensitivity SNP calls (SamTools) and one which makes high specificity SNP and indel calls (Cortex). A graph-based adjudication tool (Minos, https://github.com/iqbal-lab-org/minos) was then used to combine these results and create a final set of variants for downstream bioinformatics analysis. All strains were re-genotyped at positions that were variant in at least one CRyPTIC sample, creating a final variant call file with a call for each variant position in the *M. tuberculosis* H37Rv genome.

### Phenotyping

*In vitro* single selected mutants were further analyzed for phenotypic drug susceptibility to both BDQ and CFZ by broth microtiter dilution as a resazurin assay. All mutants were grown in antibiotic free culture media to exponential growth phase (optical density of 0.3-0.8), then diluted to the McFarland standard of 1, with an additional 1:10 dilution, 100 μl of which was seeded in 96-well flat bottom plates (about 1×10^5^ CFU per well). Next 100 μL of antibiotic was added to each well with final concentrations of BDQ as follows: 8, 4, 2, 1, 0.5, 0.25, 0.12, and 0 mg/L; and CFZ: 16, 8, 4, 2, 1, 0.5, 0.25, 0.12, and 0 mg/L. Plates were sealed with permeable tape, and incubated at 37° standing, in sealed plastic boxes. After nine days incubation at 37°, 30μl of resazurin was added to each well. After overnight incubation, fluorescence and absorbance were measure in Biotek plate reader Synergy 2. MIC was determined as the highest concentration which no bacterial growth was detected, either visually or by fluorescence measurement.

For MGIT susceptibility testing, 100μl of frozen bacterial stocks were transferred to Löwenstein-Jensen agar slants and incubated at 37° for three weeks. BACTEC™ MGIT™ 960 SIRE Kit was used and test was carried out according to manufacture instructions. Saline 0.83% solution was used for adjusting bacterial suspension concentration instead of Middlebrook 7H9 broth. Drug concentrations tested were 0.5, 1.0, and 2.0 mg/L for BDQ; 0.5, 1.0, and 2.0 mg/L for CFZ; 0.03, 0.06, and 0.12 mg/L for delamanid; and 0.5, 1.0, and 2.0 mg/L for linezolid. All MGIT tubes which were positive (growth units reached 400) before the antibiotic free growth control were considered resistant. An H37Rv WT strain was not included in this test.CRyPTIC strains were phenotyped using either the UKMYC5 plate or the updated UKMYC6 plate^83^. Plates were sealed, incubated at 37°C, and read at 14 days. In addition to manual plate readings, all plate images underwent an automated reading using AMyGDA software^46^. Plates without essential agreement for a drug MIC were marked as low quality and sent to a citizen science project (BashTheBug, http://bashthebug.net) for additional verification. Plates with exact agreement between at least two phenotyping methods were marked as high quality. Low (no method agrees) and medium (two methods with essential agreement) quality phenotypes were excluded from this analysis.

All strains included in this study had at least phenotypic data for BDQ. Some strains do not include CFZ data due to missing information, low quality analysis, or removal due to experimental error.

### *Rv0678* variant literature search

An extensive search was performed to include and summarize previously published BDQ and/or CFZ resistant associated mutations from *in vitro, in vivo*, and patient derived isolates. We used PubMed, Google, and Google Scholar to search literature published from 2014 to June 2021. Search criteria included the key “TB”, “Mycobacterium tuberculosis”, “MTB”, “bedaquiline”, “clofazimine”, “treatment”, “clinical report”, “patient”, “MDR-TB”, “XDR-TB”, “diarylquinoline”, and “drug resistance”.

Mutations in all resistance associated genes: *Rv0678, atpE, Rv1979c*, and *pepQ* were included in our final analysis, all variants with low MICs or multiple mutations in the same gene were excluded (Table 2, Table S5).

### Structural modelling

The crystal structure of Rv0678 (PDB ID: 4NB5) was visualized using UCSF Chimera^84^. Structural alignment of the wHTH domain was performed using the MatchMaker tool in Chimera with a Needleman-Wunsch algorithm using a BLOSUM-62 matrix and was iteratively pruned until no long atom-pair was > 2 Å resulting in a final average RMSD of 0.81 Å over the 5 guide structures (PDB IDs: 5HSO, 5HSM, 4FX0, 4FX4, 4YIF). All protein stability, protein-protein and protein-DNA interactions were modeled using established mCSM methods with either ligand-bound or DNA-bound Rv0678^85,86^. Mutations that presented both resistant and susceptible phenotypes were treated as resistant during statistical calculations. Screening our resistance catalogue for missense mutations that were resolved in the structure with phenotypes yielded 107 unique missense mutations. mCSM tools are available at http://biosig.unimelb.edu.au/biosig.

### Molecular dynamics simulations

All the systems simulated in the present work, including Rv0678-WT, Rv0678-A101E, Rv0678-L40V and Rv0678-L40F, were prepared using BiKi Life Sciences Software Suite version 1.3.5 of BiKi Technologies s.r.l^87^. Each simulated system consisted of Rv0678 homodimer unit X-ray structure (PDB ID 4NB5) and the mutations were generated using UCSF Chimera software^84^. The Amber14 force field was used in all molecular dynamic simulations. TIP3P waters were added to make an orthorhombic box. Adding a suitable number of counter-ions neutralized the overall system. Then, the energy of the whole system was minimized. Four consecutive equilibration steps were then performed: 1) 100 ps in the NVT ensemble at 100K with the protein backbone restrained (k=1000 kJ/mol nm^2^), 2) 100 ps in the NVT ensemble at 200K with the protein backbone similarly restrained, 3) 100 ps in the NVT ensemble at 300K with the protein backbone restrained, and 4) 1000 ps in NPT ensemble at 300K with no restraints. For atoms less than 1.1nm apart, electrostatic forces were calculated directly; for atoms further apart electrostatics were calculated using the Particle Mesh Ewald. Van der Waals forces were only calculated for atoms within 1.1 nm of one another. The temperature was held constant using the velocity rescale thermostat, which is a modification of the Berendsen’s coupling algorithm. Finally, simulations 100 ns long in the NPT ensemble at 300K were performed for each system. To detect allosteric signal transmission networks across the protein surface, defined as interconnected pocket motions, we carried out the allosteric communication network analysis using the Pocketron module in BiKi Life Sciences Suite version 1.3.5^87^.

## Supporting information

Supplemental Figures & Author List

## CRyPTIC Ethics statements as at 22 Feb 2021

Approval for CRyPTIC study was obtained by Taiwan Centers for Disease Control IRB No. 106209, University of KwaZulu Natal Biomedical Research Ethics Committee (UKZN BREC) (reference BE022/13) and University of Liverpool Central University Research Ethics Committees (reference 2286), Institutional Research Ethics Committee (IREC) of The Foundation for Medical Research, Mumbai (Ref nos. FMR/IEC/TB/01a/2015 and FMR/IEC/TB/01b/2015), Institutional Review Board of P.D. Hinduja Hospital and Medical Research Centre, Mumbai (Ref no. 915-15-CR [MRC]), scientific committee of the Adolfo Lutz Institute (CTC-IAL 47-J / 2017) and in the Ethics Committee (CAAE: 81452517.1.0000.0059) and Ethics Committee review by Universidad Peruana Cayetano Heredia (Lima, Peru) and LSHTM (London, UK).

## Authors’ contributions

LS, JC, AS, ZI, MH, KM, CU, DS, CR, KN, PF, MM and SN conceived the idea, designed the study, and analyzed and interpreted the data. All authors contributed to obtaining and assembling the data. LS, JC, AS, ZI, MM, PF and SN wrote the initial draft of the paper. All authors contributed to data interpretation, final drafting of the paper and approved the final version of the manuscript.

## Acknowledgments

We thank Dr. Musco Giovanna and Dr. Ballabio Federico (San Raffale Hospital) for their help in the molecular dynamics set up. We would like to acknowledge the technical support at Research Center Borstel, especially Vanessa Mohr, Tanja Niemann, Carina Hahn, Silvia Mass, and Doreen Beyer for their aid in the laboratory.

