## Supplemental Figures & Author List for "Deciphering Bedaquiline and Clofazimine Resistance in Tuberculosis: An Evolutionary Medicine Approach"

Supplementary Material

### Authors and members of the Comprehensive Resistance Prediction for Tuberculosis: an International Consortium

Derrick W Crook, Timothy EA Peto, A Sarah Walker, Sarah J Hoosdally, Ana L Gibertoni Cruz, Joshua Carter, Alice Brankin, Sarah Earle, Samaneh Kouchaki, Alexander S Lachapelle, Yang Yang, Timothy M Walker, Philip W Fowler, Daniel Wilson and David A Clifton, University of Oxford;

Zamin Iqbal, Martin Hunt, Kerri M Malone, Penelope Wintringer, Brice Letcher and Jeff Knaggs, European Bioinformatics Institute;

Daniela M Cirillo, Emanuele Borroni, Simone Battaglia, Arash Ghodousi, Andrea Spitaleri and Andrea Cabibbe, Emerging Bacterial Pathogens Unit, IRCCS San Raffaele Scientific Institute, Milan;

Sabira Tahseen, National Tuberculosis Control Program Pakistan, Islamabad;

Kayzad Nilgiriwala, Nerges Mistry and Sanchi Shah, The Foundation for Medical Research, Mumbai;

Camilla Rodrigues, Priti Kambli, Utkarsha Surve and Rukhsar Khot, P.D. Hinduja National Hospital and Medical Research Centre, Mumbai;

Stefan Niemann, Thomas Kohl and Matthias Merker, Research Center Borstel;

Harald Hoffmann, Katharina Todt and Sara Plesnik, Institute of Microbiology & Laboratory Medicine, IML red, Gauting;

Nazir Ismail, Shaheed Vally Omar, Lavania Joseph Dumisani Ngcamu, Nana Okozi and Shen Yuan Yao, National Institute for Communicable Diseases, Johannesburg;

Guy Thwaites, Thuong Nguyen Thuy Thuong, Nhung Hoang Ngoc and Vijay Srinivasan, Oxford University Clinical Research Unit, Ho Chi Minh City;

David Moore, Jorge Coronel and Walter Solano, London School of Hygiene and Tropical Medicine and Universidad Peruana Cayetano Heredia, Lima;

George F Gao, Guangxue He, Yanlin Zhao, Aijing Ma and Chunfa Liu, China CDC, Beijing;

Baoli Zhu, Institute of Microbiology, CAS, Beijing;

Ian Laurenson and Pauline Claxton, Scottish Mycobacteria Reference Laboratory, Edinburgh;

Robert J Wilkinson, University of Cape Town, Imperial College London and Francis Crick Institute;

Anastasia Koch, University of Cape Town;

Ajit Lalvani, Imperial College London;

James Posey, CDC Atlanta;

Jennifer Gardy, University of British Columbia;

Jim Werngren, Public Health Agency of Sweden;

Nicholas Paton, National University of Singapore;

Ruwen Jou, Mei-Hua Wu, Yu-Xin Xiao, CDC Taiwan;

Lucilaine Ferrazoli, Rosangela Siqueira de Oliveira, Juliana Maira Watanabe Pinhata, Institute Adolfo Lutz, São Paulo;

James Millard, Africa Health Research Institute, Durban;

Rob Warren, University of Stellenbosch, Cape town;

Annelies Van Rie, University of Antwerp;

Simon Grandjean Lapierre, Marie-Sylvianne Rabodoarivelo and Niaina Rakotosamimanana, Institut Pasteur de Madagascar;

Camus Nimmo, University College London;

Kimberlee Musser and Vincent Escuyer, Wadsworth Center, New York;

Ted Cohen, Yale University

Table S1: Rv0678 variant diversity in *in vitro* population. Total variant diversity in resistance associated genes was assessed thought collection of all mutant colonies which grew on selective agar plates. Mutant populations were collected per experimental condition (drug exposure, concentration, and isolates timepoint) and deep sequencing was performed (whole genome sequencing). Substitution annotated by coding position, and “alternative allele” describes the variant at the base pair position. The highest frequency in which the substitution was found in any given population and number of reads with a phred score greater than 20 were included. “Isolated from” denotes whether the substitution was observed after BDQ or CFZ (or both) exposure. It is noted if the substitution was also selected in as a single clone (Table 1).

BDQ – bedaquiline, CFZ – clofazimine, del – deletion, fs – frameshift, ins – insertion, NA – not available, SNP– single nucleotide polymorphism, * – stop codon insertion

Table S2: MIC verification by MGIT. Additional drug susceptibly testing was carried out in MGIT BD system with mutant clones selected after evolutionary experiments. R or “resistant”, indicates the alone grew at correlating concentration. The WHO defines bedaquiline resistance in MGIT as 1.0 mg/mL, clofazimine as 1.0 mg/L, delamanid as 0.06 mg/L, and linezolid as 1.0 mg/L.

Table S3: Secondary mutations co-selected with resistant variant in vitro. Secondary mutations are off-target variants which were co-selected with a resistant variant. All mutations described were detected by whole genome sequencing, aligned to the reference strain, and compared to the wild type ancestor genome. The number of times the variant was selected under “# of isolates”.

BDQ – bedaquiline, CFZ – clofazimine, del – deletion, fs – frameshift, ins – insertion, Substi. – substitution, * – stop codon insertion

Table S4: Patient isolates collected by CRyPTIC. All isolates which were included in this study that were collected by CRyPTIC (not previously published). A total of 235 isolates were included which harbored a mutation in *Rv0678*, *atpE*, *Rv1979c*, *pepQ*, and were phenotypically verified on UKMYC5/6 plates, for at least bedaquiline (also most with clofazimine).

BDQ - bedaquiline, CFZ - clofazimine, del – deletion, DST-drug susceptibility testing, ins – insertion, LAM – Latin American, SNP – single nucleotide polymorphism, * – stop codon insertion

Table S5: Reference catalogue of variants implicated in bedaquiline and/or clofazimine resistance associated genes, *Rv0678*, *atpE*, *pepQ*, and *Rv1979c*. Catalogue include variants described in *in vitro*, *in vivo*, and patient isolates from our *in vitro* and CRyPTIC datasets plus previously pushed literature. Variant inclusion from literature were filtered for resistance and borderline MIC increases only and no more than one mutation in a resistance associated genes. Drug susceptibility testing method and critical concentrations (mg/L) were:

[A] BD-MGIT: 1.0 BDQ & CFZ; [B] UKMYC5/6: 0.25 BDQ & CFZ [D] Alamar blue/resazurin assay, [E] 7H10 plates, [F] 7H11 plates.

Phenotype interpretation was based off of: [A] 1mg/L BDQ/CFZ^80^; [B] 0.12-0.25mg/L BDQ/CFZ^75^, [C] defined by study authors, [D] 0.5mg/L BDQ^81^/CFZ^82^, [E] 0.25 mg/L BDQ^80^ -CFZ determined by study authors

del – deletion, fs – frameshift, ins – insertion, NA – not available, SNP – single nucleotide polymorphism, *– stop codon insertion

Table S6: Predicted biophysical effects of Rv0678 mutations. mCSM predicted effects on protein stability (DDG, kcal/mol), dimer stability, dimer stability of the DNA-bound conformation, and DNA-binding affinity. Distance to the interface (Å) and residue solvent accessible surface area was calculated for the non-DNA bound conformation of Rv0678. Significant effects on stability were defined as >1kcal/mol.

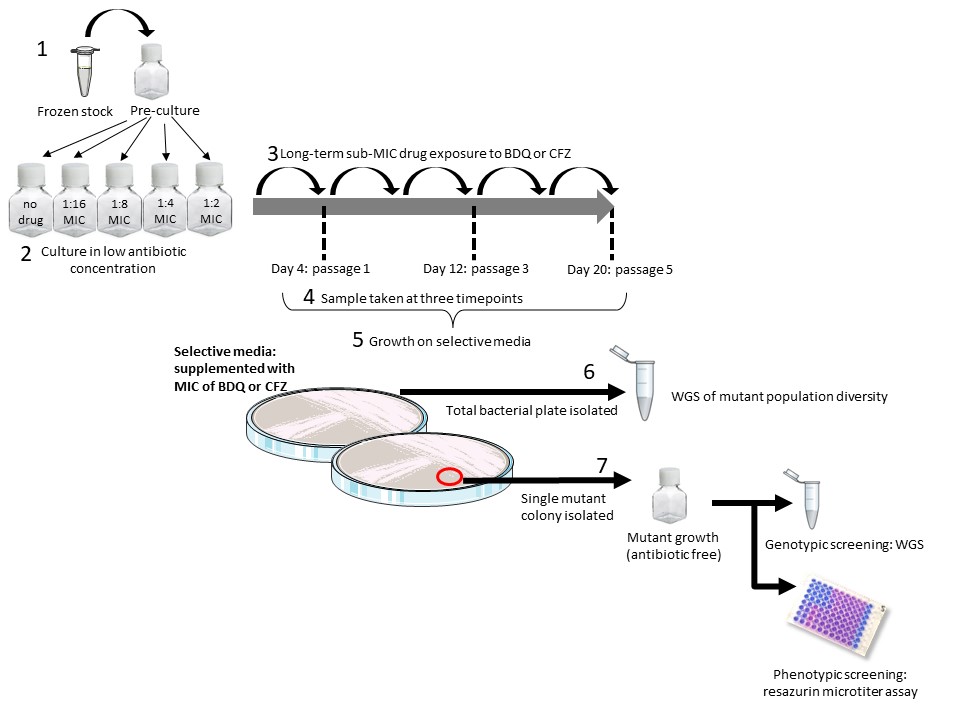

Figure S1: *In vitro* evolutionary experimental design for resistant variant selection, detection and analysis. (1) First, a pre-culture was started from frozen stocks of *Mycobacterium tuberculosis* complex strain H37Rv, (2) at exponential growth phase bacteria were transferred into new culture bottles and exposed to sub-minimum inhibitory concentrations (MIC) of antibiotics. (3) Bacteria were culture for 20 days including five bacterial passages, (4) cultures were sampled at passages 1, 3, and 5, (5) and grown on selective media plates, supplemented with the MIC of the antibiotic (0.12-0.25 mg/L bedaquiline or 0.25 mg/L clofazimine). (6) After growth on selective media all colonies were pooled and deep sequencing of the heterogenous population was analyzed. (7) Single mutant colonies also were isolated from selective media plates and characterized by whole genome sequencing and MIC tested by resazurin microtiter plate assays.

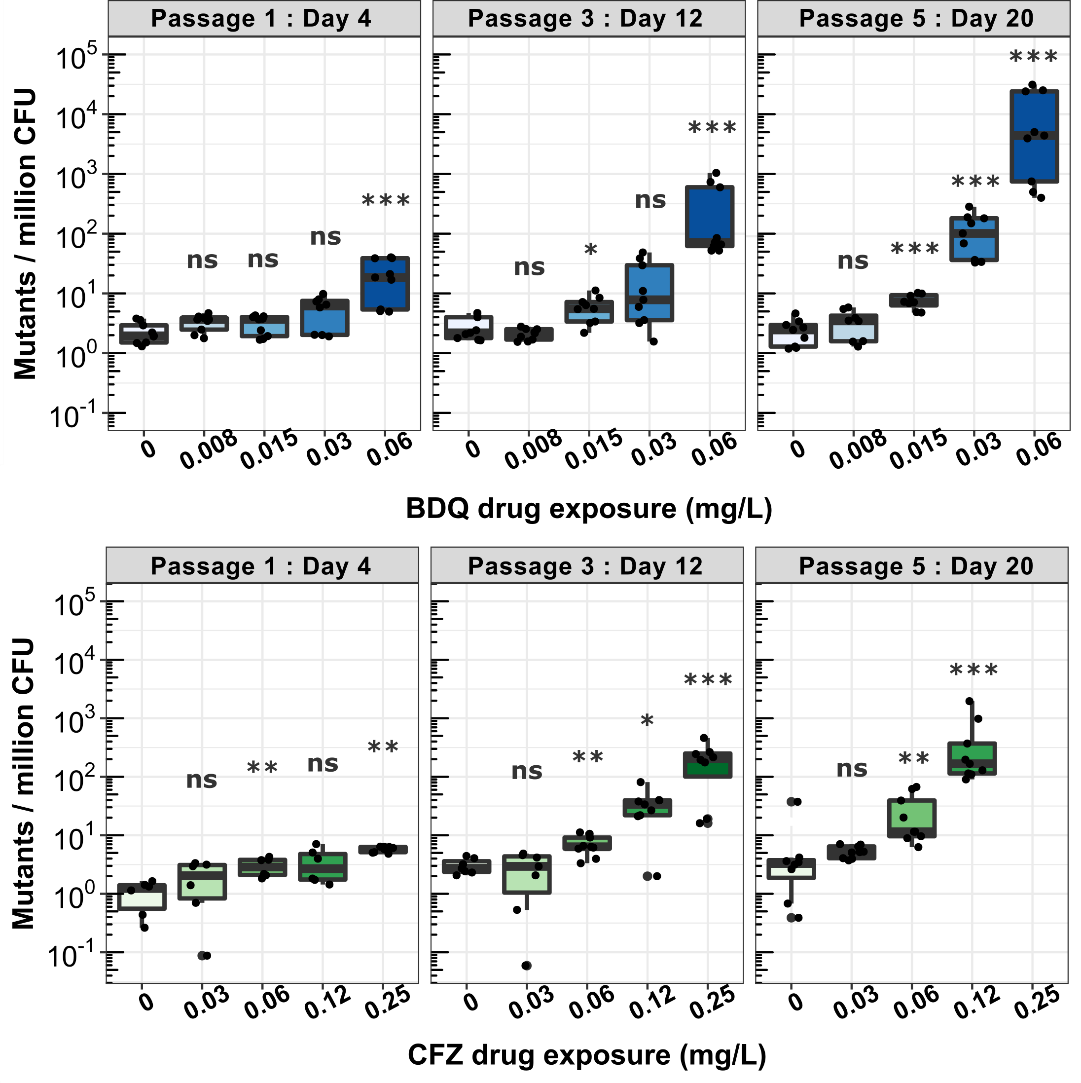

Figure S2: Sub-lethal exposure of antibiotics enriches drug resistant populations in a dose dependent manor. The *Mycobacterium tuberculosis* complex lab strain H37Rv was exposed to four concentrations of bedaquiline (BDQ) (0.06, 0.03, 0.015, and 0.008 mg/L) or clofazimine (0.25, 0.12, 0.06, and 0.03 mg/L), plus an antibiotic free control (0 mg/L BDQ/CFZ), with the highest concentration at 1:2 the minimum inhibitory concentration (MIC), MIC= 0.12mg/L for BDQ and 0.5mg/L for CFZ. The bacteria were exposed to the antibiotic for 20 days, consisting of 5 culture passages. Bacterial samples were evaluated at three timepoints during the experiment, after day 4, day 12, and day 20. Cultures were diluted and plated on 7H11 agar plates, supplemented with and without the MIC of each drug. After 14 to 21 days of growth, colony forming units (CFU) were counted. Mutants per million CFU was calculated by dividing number of mutant CFU/mL by total CFU/mL, then multiplied by 10^6^.

Statistics: Three independent experiments were conducted, with 1 biological replicates per experiment, and 3 to 5 technical replicates per biological replicate (9 to 15 values). Statistics was calculated as nonparametric multiple contrast test (Kruskal) with a confidence interval of 95%, p-values between drug exposed and the antibiotic free control (0 mg/L BDQ): *p<0.05, **p<0.01, ***p<0.001

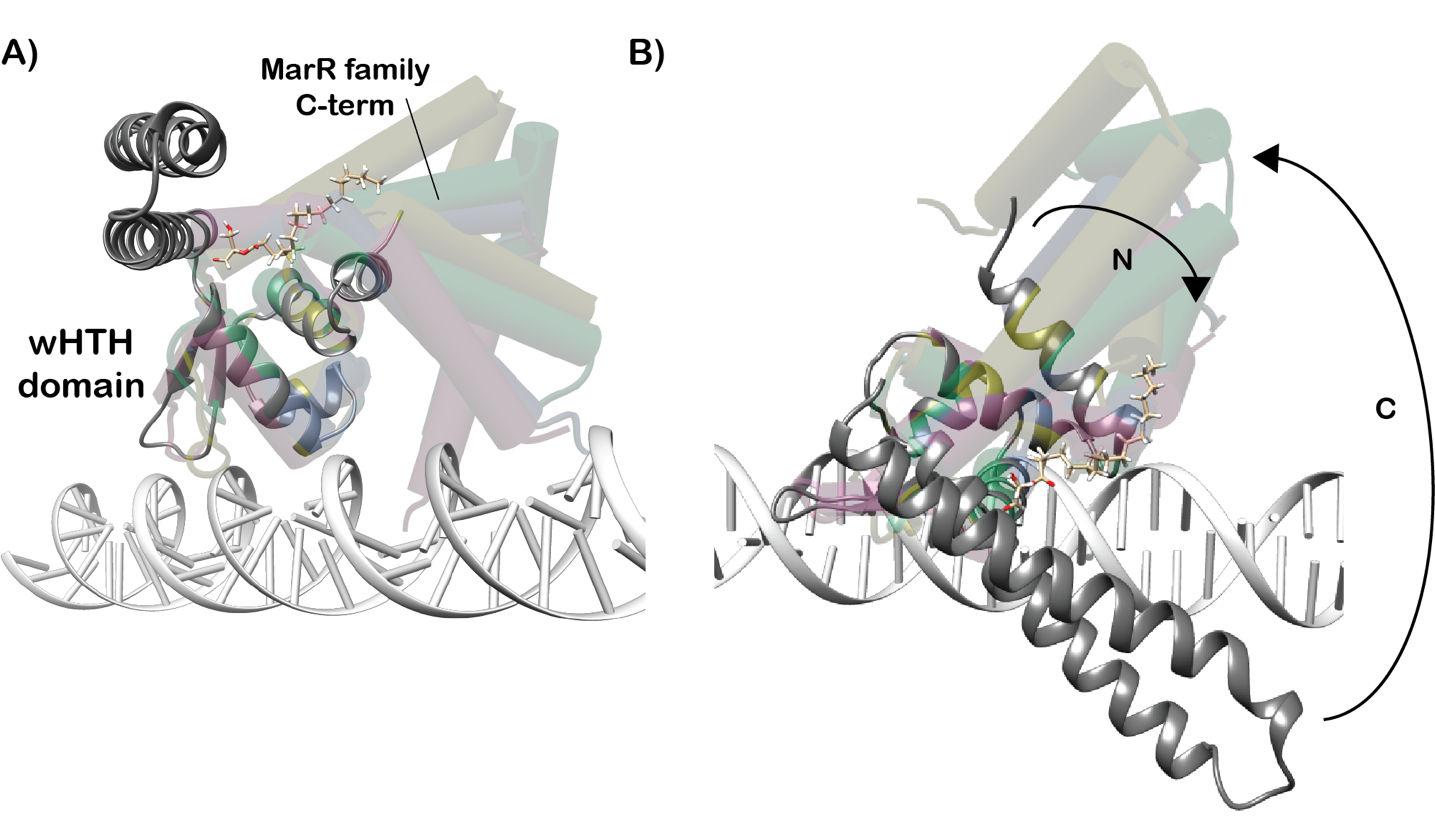

Figure S3: Rv0678 C-terminal domain undergoes conformational change upon DNA binding. (A) Structural alignment of Rv0678 with 5 other MarR-family proteins from Mtbc. Average RMSD across 5 alignments is 0.81 Å. (B) The C-terminal domain conformational change likely involves concerted and counter-directional rotations of the N-terminal helix and C-terminal dimerization domain.

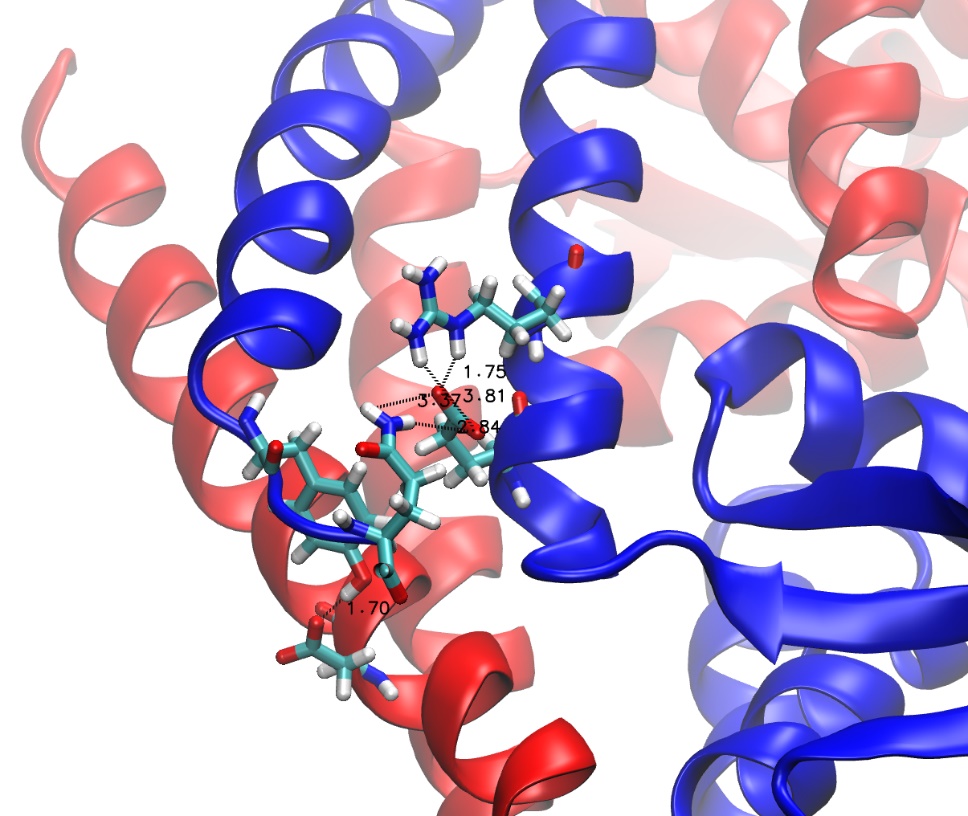

Figure S4. Zoom-in of the persistent interaction along the A101E simulations. The two Rv0678 are shown in red and blue cartoon. The residues involved in the interaction are shown in licorice. In dashed lines are shown the interaction between residues along with the distance in Angstroms unit.

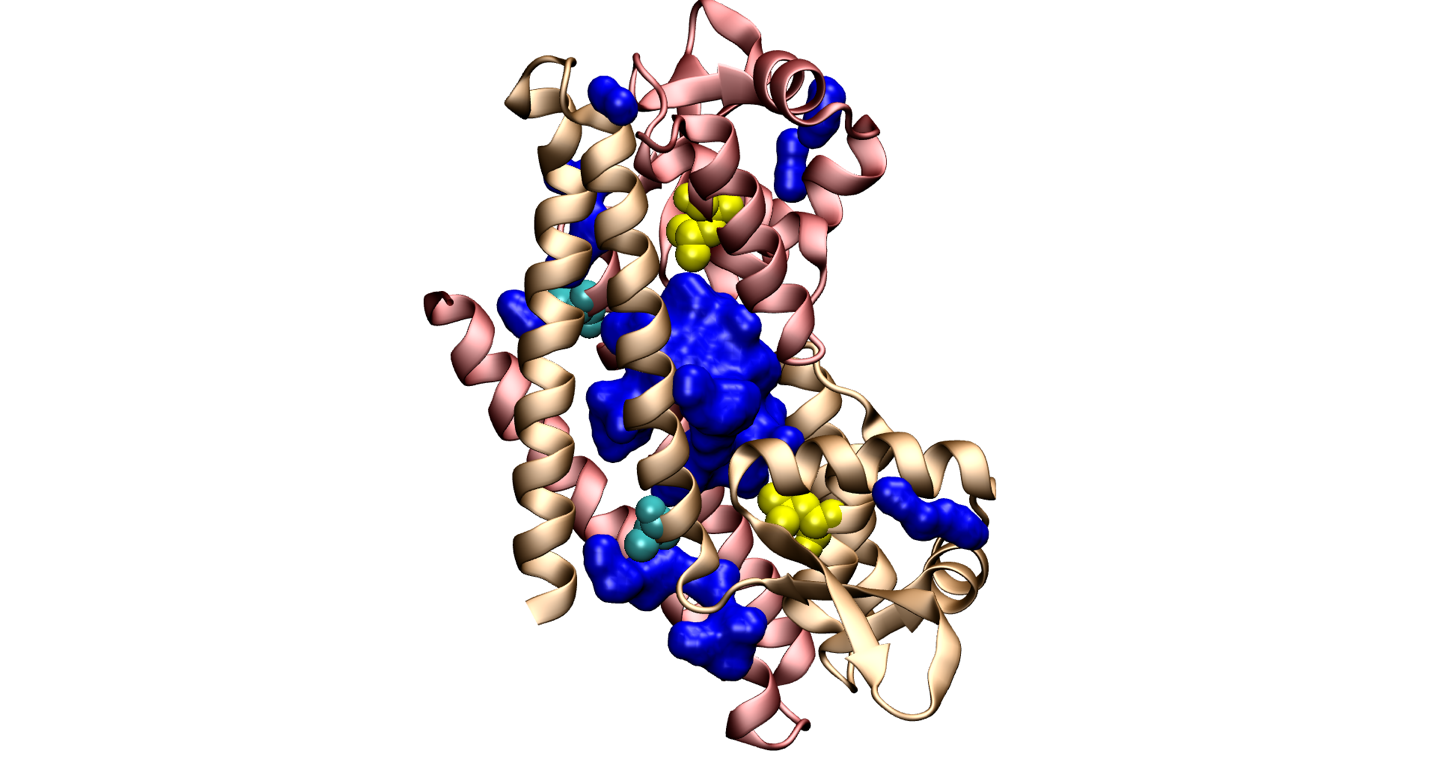

Figure S5: Localization of the main pockets in Rv0678 (4NB5), shown in blue surface. The two monomers are shown in pink and orange cartoon. The residues A101 and L40 are shown in cyan and yellow sphere vdW respectively.
